# Reduced belief updating impairs adaptive step initiation in older adults

**DOI:** 10.64898/2025.12.19.695189

**Authors:** Alan Bince Jacob, Sonja A. Kotz, Ryszard Auksztulewicz, Markus Köpcke, Kenneth Meijer, Tjeerd W. Boonstra

## Abstract

Ageing is associated with declines in mobility and balance that threaten independence and increase fall risk. Although these changes are often attributed to deterioration of muscular and sensory systems, growing evidence suggests that age-related alterations in cognitive control also contribute to postural instability and gait impairments. Here, we examined whether shifts in cognitive control strategy underlie impaired gait initiation in older adults by modelling how individuals update beliefs about upcoming actions. Older and younger adults performed a choice-stepping go/no-go task in which step likelihood was manipulated across trials. We measured anticipatory postural adjustments (APAs), the preparatory weight shifts preceding stepping and modelled age-related differences in APA onset using a hierarchical Bayesian framework. Older adults initiated slower APAs, particularly after repeated go trials. Computational modelling revealed reduced learning rates and heightened sensitivity to uncertainty in older adults, identifying altered belief updating as a cognitive mechanism shaping age-related changes in adaptive stepping.

## 1. Introduction

Postural control deteriorates with age, resulting in increased postural sway, diminished coordination, and reduced adaptability to changing environmental conditions. These changes increase falling risk, limit mobility, and significantly affect the quality of life in older adults ^1^. Reasons for age-related postural control decline are multifactorial, and both peripheral and central processes contribute to diminished balance. Peripheral processes include age-related deterioration of muscle strength ^2,3^ and weaker force generation with ageing ^4^. Age-related changes in the central nervous system include increased sensorimotor processing delays ^5^, secondary task deficits^6^ and increased attentional demands ^7^. While it is well established that both central and peripheral factors contribute to postural instability ^1^, their relative contributions are not fully understood.

Evidence for age-related decline in cognitive control of gait comes from dual-task paradigms and neuroimaging studies of gait, which showed increased cognitive-motor interference and greater activation of prefrontal cortical regions functionally associated with postural control in older adults^8,9^. These changes engage various cognitive subfunctions, including attention, planning, response monitoring, response inhibition, and volition ^10^. Neuroimaging studies have shown that executive control regions such as dorsolateral prefrontal cortex, which support attention and postural regulation, exhibit age-related decline in grey matter volume associated with slower information processing and alterations in spatial and temporal gait characteristics^11,12^. Braver and colleagues proposed that cognitive control operates via two distinct modes: proactive control, which supports anticipatory and sustained adjustments, and reactive control, which is engaged only after the detection of conflict between expected and perceived information^13,14^. Variations in cognitive control across conditions and individuals can therefore be understood as shifts in control strategy that optimize information processing given task demands and individual characteristics. Reduced adaptability to environmental changes and diminished stability during gait with ageing may reflect such a shift in control strategy.

Incorporating neurocognitive paradigms into a postural task – rather than adding them as a separate secondary task – may help to disentangle these cognitive subfunctions. For example, by adopting different stepping conditions with contrasting balance demands, more anticipatory or reflexive stepping strategies could be induced. This approach revealed distinct cortical activity between stimulus appearance and response, mainly increased theta/alpha power in the supplementary motor area (SMA) and alpha/beta suppression in the primary motor and parietal cortices^15^. Similarly, an experimental paradigm using a modified Stroop-like design to manipulate cognitive conflict during step initiation^16^ revealed worse inhibitory control and reduced cortical activation in the SMA and dorsolateral prefrontal cortex in older participants.

By manipulating stimulus-stimulus contingency in a go/no-go design, Ziri and colleagues^17^ dissociated proactive and reactive inhibition processes during gait initiation, demonstrating how preparatory inhibition engages occipito-parietal and motor cortical areas, while stimulus-driven reactive inhibition involves frontal and motor regions. More broadly, serial dependencies between stimuli have been shown to influence behavioural outcomes across perceptual and cognitive tasks^18,19^. Manipulating these dependencies provides a powerful approach to probe higher-order cognitive control functions^20,21^.

In this context, the theory of predictive coding offers a unifying framework, describing a continuous hierarchical process whereby the brain updates its initial beliefs (priors) by minimizing prediction errors to enable flexible adaptations ^22–24^. Recent work in statistical learning has demonstrated how this framework can be used to assess how top-down expectations are influenced by statistical regularities across perceptual, abstract, and motor levels – thereby allowing stream of information from successive stimuli to improve perceptual stability for guiding response selection^25^. Connecting this framework with dual-mechanisms of control, proactive control can be understood as the sustained stabilization of probabilistic priors supporting anticipation, and reactive control as stimulus-driven updating of these expectations, with their relative influence determined by Bayesian precision weighting^26^. With ageing, there might be a shift from a learning-focused mode to more rigid predictions, leading to a reliance on well-established probabilistic priors rather than rapid belief updating of probabilistic beliefs^27^. To assess how this shift affects the relative precision weighting of anticipatory and reactive modes, the Hierarchical Gaussian Filter (HGF)^28,29^ – a computational model of belief updating inspired by predictive coding framework – can be applied to the control of posture and gait. This approach enables the investigation of age-related differences in reliance on probabilistic priors ^30,31^ and diminished stimulus-driven updating of internal beliefs, as reflected by perceptual learning rate changes within a dynamic environment ^32^.

In this study, we tested younger and older adults while they performed a choice stepping task with a go/no-go paradigm in which we experimentally manipulated the odds ratio between go and no-go trials (3:1 and 1:3). We measured both the foot lift-off reaction times (RTs) and anticipatory postural onsets and assessed the serial dependencies of stimuli and RTs using the HGF. We hypothesized that older adults would show slower anticipatory postural onsets and reduced learning rates, indicating reduced belief updates, increased reliance on prior expectations, and lower dependence on sensory evidence (stimuli). Through this Bayesian approach of learning in an uncertain environment, we show mechanistically how older adults shift their control strategy towards reactive control. This adaptation minimizes the heightened risks of prediction errors under environmental constraints, thereby sustaining stability and performance in postural control and gait.

## 2. Results

### 2.1. Participant Characteristics

In total, 24 younger and 25 older adults participated in the study (Table 1). As expected, older participants were slower than younger participants on the timed-up and go (TUGT; *7.7* ± *1.2 s, range = 6.0 - 11.0 versus 5.0* ± *0.8s, range = 3.7 - 7.0*; *p < .001*). All participants completed the TUGT below the cutoff performance time of 13.5 seconds ^33^ and older participants stayed within the normative reference range between 8 to 11 seconds ^34^ for healthy adults aged 65 to 85 years, indicating normal functional mobility and low fall risks. BBS scores (*M = 55.4* ± *0.9, range = 53.0 - 56.0*) indicated that older adults in this study were not at risk of falling and showed good functional balance as they scored close to the maximum value of 56 and their scores were well above the BBS cut-off of 46 ^35^.

**Table 1.**
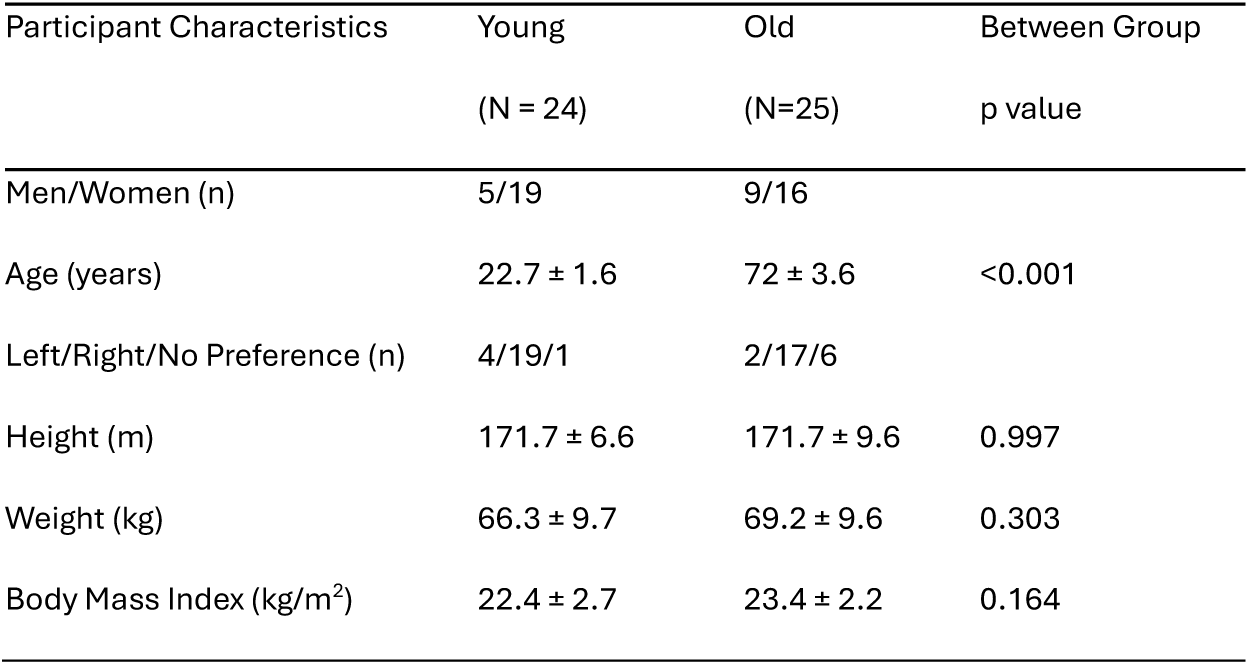
Participant Summary Table (mean ± *SD*)

| Participant Characteristics | Young<br>(N = 24) | Old<br>(N=25) | Between Group<br>p value |
| --- | --- | --- | --- |
| Men/Women (n) | 5/19 | 9/16 |  |
| Age (years) | $22.7 \pm 1.6$ | $72 \pm 3.6$ | <0.001 |
| Left/Right/No Preference (n) | 4/19/1 | 2/17/6 |  |
| Height (m) | $171.7 \pm 6.6$ | $171.7 \pm 9.6$ | 0.997 |
| Weight (kg) | $66.3 \pm 9.7$ | $69.2 \pm 9.6$ | 0.303 |
| Body Mass Index (kg/m<sup>2</sup>) | $22.4 \pm 2.7$ | $23.4 \pm 2.2$ | 0.164 |

### 2.2. Anticipatory Postural Onset Time

We determined the occurrence of anticipatory postural adjustments (APAs) in step initiation after the presentation of a go cue in a choice stepping task (Fig 1A, B). An APA is the force exerted by the stepping leg to shift the weight onto the stance leg prior to lift-off. Younger participants showed an APA in 97% of the go trials and older participants in 95% of the go trials.

**Fig. 1.**
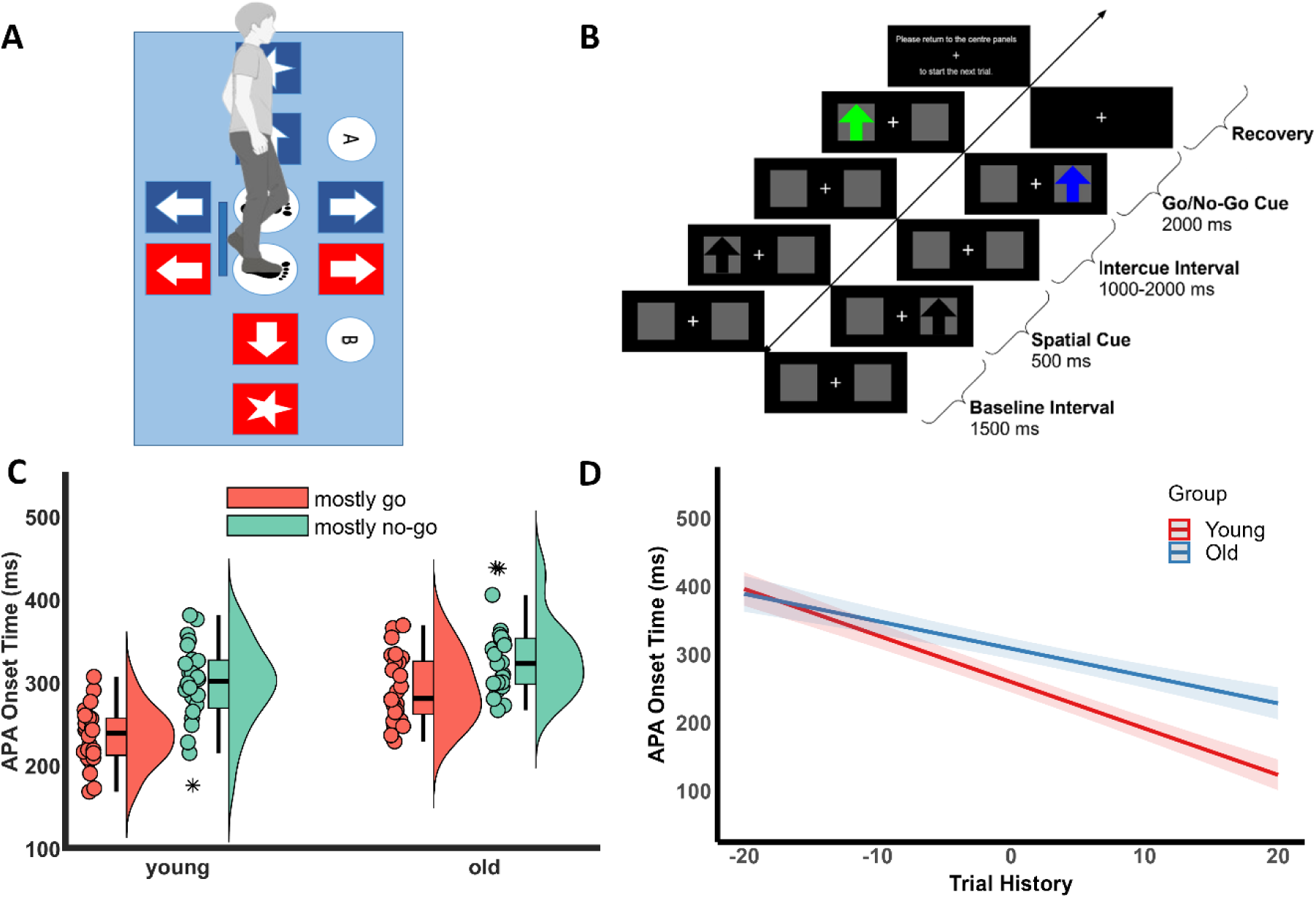
Experimental setup of the choice stepping task. **(A)** The participant stood on the middle panels of the computerized stepping mat in front of a computer monitor. The monitor displayed instructions to perform the choice stepping task. The participant was instructed to step onto the forward panel if a green arrow (go) is displayed and remain on the centre panels in case of a blue arrow (no-go). The participants stepped with either the left or right foot depending on the position of the green arrow. **(B)** The time courses of events for a go-left trial and a no-go-right trial, respectively. The trial starts with a fixation cross displayed in the centre of the monitor (1500 ms), followed by a spatial cue (500 ms). A black arrow is displayed in either the left or right panel for stepping with the left or right foot, respectively. After an inter-cue interval (1000-2000 ms), a go/no-go cue is presented (2000 ms) indicating either go (green arrow) or no-go (blue arrow). If the go cue is presented, the participant makes a step forward. At the end of the trial, an instruction is displayed to return to the centre panels. **Anticipatory Postural Onset Time (C)** Raincloud plot and boxplot with median line for mean APA onset time of young and old participants. The scatter plot indicates individual mean APA onset time, colours indicate conditions, the box indicates interquartile range (IQR) between 25^th^ to 75^th^ percentiles (median line in the 50^th^ percentile), and the whiskers represent most extreme datapoints in each group within 1.5 times the IQR below the 25^th^ percentile and above the 75^th^ percentile respectively. **(D)** Marginal effects of the interaction between trial history and group. The slope lines with 95% confidence intervals in the plot show how APA onset time changes as a function of trial history in young (red) and old (blue). Here the negative numbers in trial history indicate consecutive no-go and positive numbers indicate consecutive go.

Comparing the vertical force across groups (Fig S1.1A), older adults showed higher normalized force at maximum APA (0.7) than younger adults (0.66), but this numerical difference was not statistically significant (*p = 0.246*). Despite absent group differences (Table S1.1), a main effect of condition and an interaction effect of condition x group (*F (1, 47.9) = 10.9, p = 0.002, partial η² = 0.19, 90 % CI [0.05,0.34]*) was observed. Although the effect size of the interaction was small to medium, post-hoc analysis showed a small force difference for younger adults between the mostly-no-go and mostly-go conditions (Fig S1.1B).

There were no false alarms and miss rate was lower in younger adults than older adults (*0.01* ± *0.02 versus 0.04* ± *0.04; p = 0.001*). We then compared RTs across groups and conditions using a LMM for touchdown time, lift-off time, maximum APA time, and APA onset time generated from the latencies of these force profiles in the remaining go trials (Fig. S2.1). We used on average 116.2 ± 5.8 trials for young and 76.2 ± 4.3 trials for old participants in the analyses (Table S1). As we found qualitatively similar effects at all four latencies (Table S2.1), we focus on APA onset in the remainder of this paper.

For APA onset time (Fig 1C), we found average faster APA onset times were observed in mostly-go compared to mostly no-go, and younger adults had faster APA onset time than older adults (Table S2.1). A significant interaction effect (*F (1, 46.5) = 7, p = 0.011, partial η² = 0.13, 90% CI [0.02,0.28]*) between condition and group was also found. The effect size of the interaction was medium to large. Post-hoc analysis (Table S2.2) revealed significant differences in three pairs: younger adults had faster APA onset time than older adults in mostly go, and both younger and older adults were faster in mostly go than mostly no-go condition.

LMM was also conducted on the movement time (time between foot lift-off and touchdown), but no significant interaction effects were observed (Table S2.1).

### 2.3. Trial History

The previous analysis compared average RTs in the mostly go and mostly no-go conditions. To assess serial dependencies in RTs, we then performed a LMM on APA onset time as a function of trial history (Table S2.3). Here, APA onset time showed a main effect of trial history, highlighting that, APA onset time decreased with consecutive go trials and increased in go trials after consecutive no-go trials. This main effect suggests robust serial dependency on average across participants. Looking at individual participants, we observed negative slopes in 92% of our total participants (100% in young and 84% in old; Fig S2.2 and S2.3), showing a decrease in APA onset time with trial history. A trail history × group interaction (*F (1, 43.4) = 7.7, p = 0.008, partial η² = 0.15, 90 % CI of partial η² [0.02,0.31]*) effect was also observed, indicating a more negative slope in younger adults compared to older adults (Fig 1D). This decrease in slope with trial history suggests that younger adults had faster APA onset time than older adults for consecutive go trials compared to consecutive no-go trials. From these effects, we observed stronger evidence of serial dependency in younger adults than older adults.

### 2.4. Hierarchical Gaussian Filter

To identify individual differences in anticipation, learning, and decision making that underlie the age-related effects of serial dependency, we fitted the HGF to the APA onset times (Fig. 2A), using the prior values for response model parameters described in Section S3. By fitting the model to the empirical data of each participant, we obtained trial-by-trial posterior expectations. These belief trajectories show the inferred posterior estimates and their associate uncertainty, which evolved as a function of the sequential order of go and no-go trials (Fig. 2B). The magnitude of these belief updates is scaled by the higher-level learning rate ω. Seven parameters (ω, β_0_-β_4_, and ζ) were estimated for each participant.

**Fig. 2.**
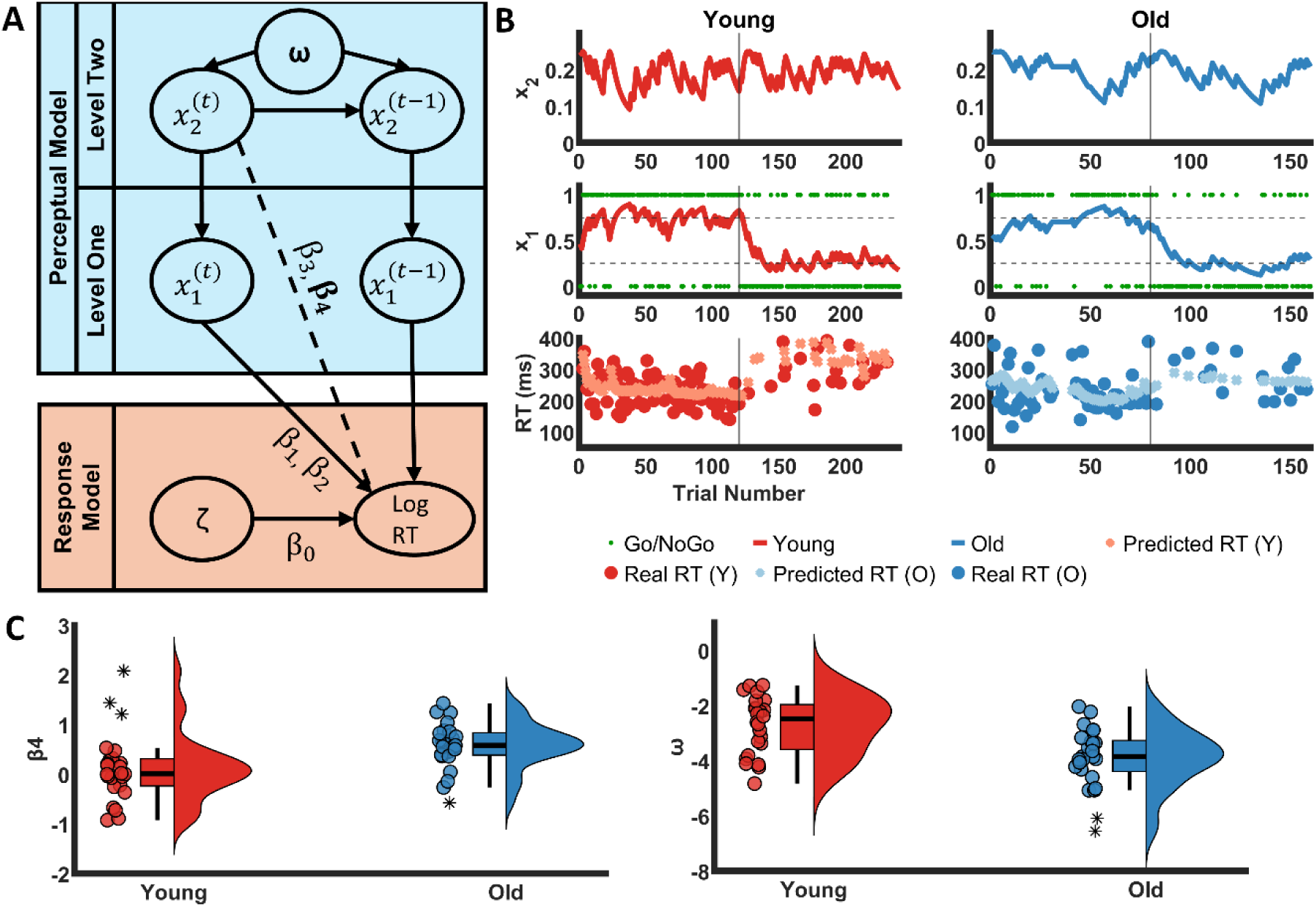
Hierarchical Gaussian Filter. **(A)** Flow diagram of HGF showing the division of two main models: the perceptual and response model. The perceptual model consists of two levels and the learning rate is derived from the second level. **(B)** Estimates of environmental uncertainty beliefs connected to β_4_, posterior expectations, and predicted vs real APA onset times in a representative young and old participant with learning rates close to the group mean. Dotted lines on y-axis indicate the odds ratio of go and no-go stimuli (0.75 and 0.25) and the solid line on x-axis indicates trial number where the shift in odds ratio occurs **(C)** Raincloud plots of β_4_ and ω (learning rate) with median and individual datapoints. The box represents interquartile range (IQR) between 25^th^ to 75^th^ percentiles (median line in the 50^th^ percentile), and the whiskers represent most extreme datapoints in each group within 1.5 times the IQR below the 25^th^ percentile and above the 75^th^ percentile respectively.

We then performed independent samples t-tests between groups to examine age-related differences in these parameters (Tables S3.1 and S3.2). Younger participants (*M = −2.7, SD =1*) showed a higher learning rate ω (*t(47) = 3.89, p = 0.002 [corrected], Cohen’s d = 1.11, 95 % CI of Cohen’s d [0.50,1.71]*) than older participants (*M = −3.9, SD = 1.1*). In contrast, older participants (*M = 0.57, SD = 0.46*) showed a larger β_4_ than younger participants (*M = 0.12, SD = 0.70*), (*t(47) = - 2.68, p = 0.03 [corrected], Cohen’s d = −0.76, 95 % CI of Cohen’s d [−1.34, −0.18]*; Fig 2C). This parameter represents how strongly unexpected uncertainty influences RTs, with higher β_4_ values indicating greater sensitivity to sudden environmental changes and consequently slower motor responses, hence slower APA onset times. The offset parameter β_0_, amounting to overall APA onset times, independent of trial-by-trial dynamics, was larger (*t(47) = −3.19, p = 0.009 [corrected], Cohen’s d = −0.91, 95 % CI of Cohen’s d [−1.50, −0.32]*) in older participants (*M = 5.6, SD = 0.3*) than younger participants (*M = 5.4, SD = 0.3*). Looking at the three parameters, we see large effect sizes in ω and β_0_, and medium to large effect size in β_4_. In β_4_, we also observe larger variability in younger participants than older participants compared to the mean (Fig 2C).

### 2.5. Posterior Predictive Checks

To check whether the observed group differences in ω and β_4_ explained the experimental effects on APA onset time that were observed in the empirical data, we performed posterior predictive checks. APA onset time was simulated for 240 trials using values of ω and β_4_ ranged within participant’s parametric space and mean RT was computed for different conditions (mostly go and mostly no-go) or for different trials histories using 100 different random sequences of go/no-go inputs. In the simulated data, we observe a general increase in RTs at higher values β_4_ across conditions. The effect of ω differed for different values of β_4_: for β_4_ = 0.1 higher values of ω resulted in lower RTs for mostly-go, while for β_4_ = 0.6 higher values of ω resulted in higher RTs for mostly-go. The differences in RTs between conditions also depended on the specific combination of β_4_ and ω, but for most combinations RTs were higher in the mostly no-go compared to the mostly-go condition. Taking the average values of β_4_ and ω in the young (β_4_ = 0.1, ω= −3) and old (β_4_ = 0.6, ω= −4) , we observe a similar pattern as in the empirical data: RTs are lower in the young group, RTs are lower in the mostly-go condition and this differences in larger in the young group (Fig. 3A). A similar pattern is observed for the differences in RTs as a functioning of trial history: There is a general decrease in RT after more subsequent go trials, and this trend is stronger for the young group (Fig. 3B). Hence, these model-based findings showed that our model adequately captured patterns of results found in the empirical data.

**Fig. 3.**
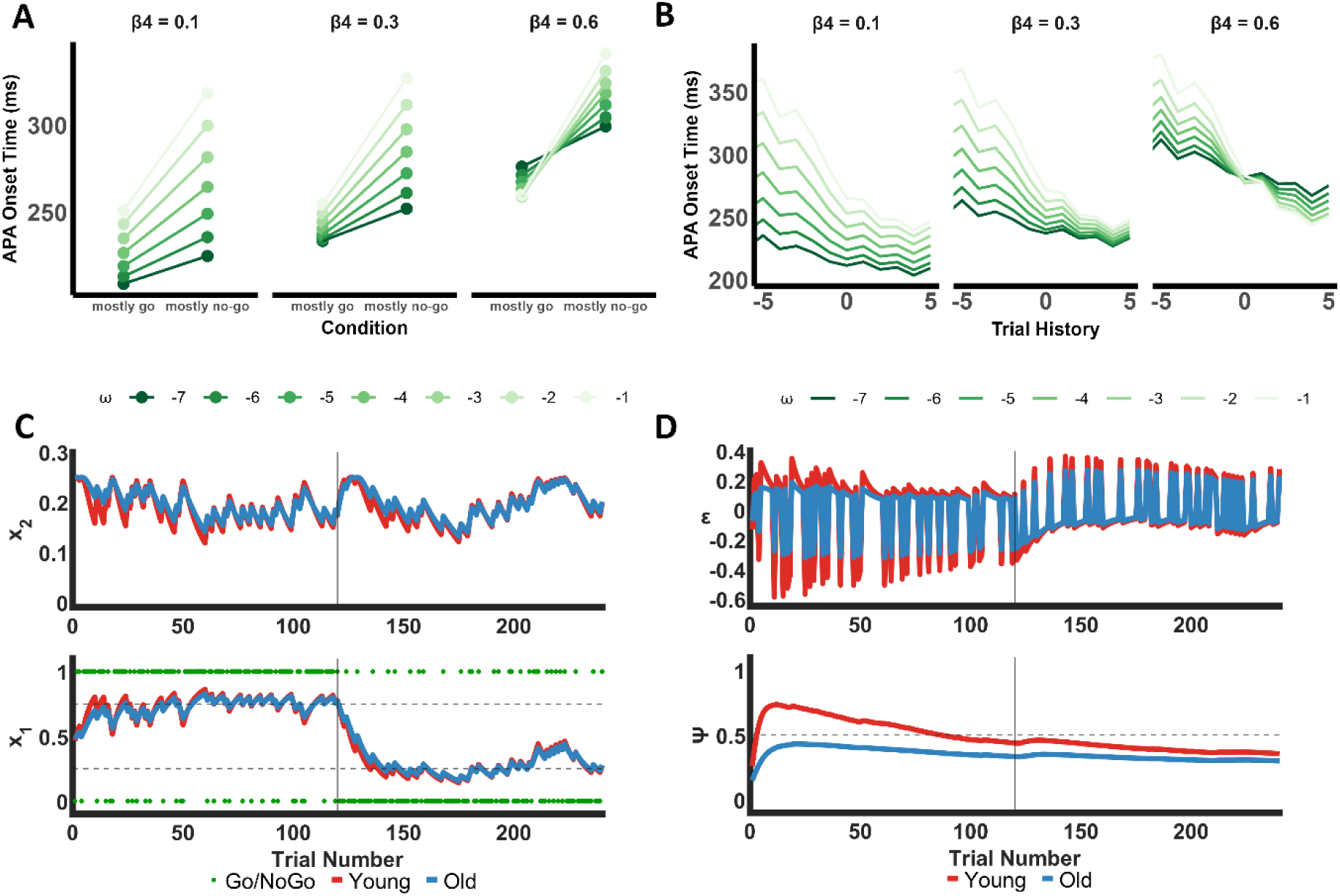
Posterior Predictive Checks and Hidden States of HGF. **(A)** Simulated mean APA onset time for mostly-go and mostly no-go conditions for a parameter sweep of ω (−7 to −1) and β_4_ (0.1 to 0.6) to evaluate the relative contributions of ω and β_4_ from HGF on the mean APA onset time in both conditions. The average parameter values in young (β_4_ = 0.1, ω= −3) and old (β_4_ = 0.6, ω= −4) revealed lower APA onset times in the mostly-go than in the mostly no-go condition, higher APA onset times in old compared to young, and a reduced differences between the two conditions in old compared to young. **(B)** Simulated mean APA onset time as a function of trial history for a parameter sweep of ω (−7 to −1) and β_4_ (0.1 to 0.6). As observed in empirical data, APA onset time decreases with trial history across different ω and β_4_. Larger values of β_4_ resulted in an overall increase in simulated RTs, while larger values of ω resulted in steeper slopes of simulated RTs with trial history. **(C)** Simulated posterior expectation (x_1_) and estimates of environmental uncertainty connected to β_4_ (x_2_) using the average parameter values of ω and β_4_ from young (red) and old (blue). The vertical solid line indicates the change point of condition (from mostly-go to mostly-no-go) and the horizontal dashed lines indicate expectancy of go-cue - 0.25 (25% go) and 0.75 (75% go). The green dots indicate binary inputs for no-go cue (0) and go cue (1) **(D)** Simulated precision-weighted prediction errors (ε) and precision weights (ψ) of prediction errors across trials in young (red) and old (blue). The horizontal dotted line indicates ψ = 0.5 and the vertical solid lines indicate the change point of condition (from mostly-go to mostly-no-go).

### 2.6. Latent variables of HGF

To take a closer look at how ω and β_4_ capture differences in age-related stimulus-driven learning across trials, it was necessary to look at the trial-by-trial dynamics of precision-weighting on prediction errors with age. To do this, we simulated hidden states of the HGF’s perceptual model by using the mean of the fitted ω and β_4_. In the perceptual model, the simulations showed an increased variability in posterior expectation (x_1_) in mostly-go than in mostly no-go. Consistent with model fitting results (Section 2.4), belief trajectories simulated using the average parameter values of ω and β_4_ from young and old showed more dynamic updating in young participants (Fig 3C). When the probability of go cue was high (75% go probability), young exhibited higher posterior expectations than old. Conversely, in mostly no-go condition (25% go probability), their posterior expectations were lower, reflecting more accurate tracking of environmental contingencies. Younger adults also demonstrated stronger belief updates regarding environmental uncertainty (x_2_), indicating more flexible learning. This shows how the perceptual model captured the dynamic learning behaviour of young and lower trial history dependent behaviour of old as younger adults made use of odds ratio of stimuli and changing environmental conditions to anticipate trial wise responses. The latent variables of the perceptual model in the HGF revealed larger precision weighting on prediction errors (ψ) in young than old (Fig 3D). This pattern was consistent regardless of condition order, i.e., whether the participants started with mostly-go or mostly no-go trials (Fig S3.1). Correspondingly, younger adults exhibited larger precision-weighted prediction errors (ε), indicating that they placed greater emphasis on prediction errors to improve their belief updates based on sensory inputs for faster initiation of stepping response in anticipation of a go/no-go cue.

## 3. Discussion

We used a choice-stepping task to test whether older adults rely more heavily on prior beliefs and exhibit reduced flexibility in adapting to environmental changes. By manipulating the ratio of go to no-go trials and modelling serial dependencies in anticipatory postural adjustment (APA) onset times, we identified several age-related differences in the cognitive control of step initiation. First, although older adults responded more slowly than younger adults, this difference was most pronounced during sequences of consecutive go trials. Second, these effects emerged early, at the level of APA onset (∼200 ms after cue onset). Third, computational modelling of the empirical data showed that these differences were captured by a lower learning rate (ω), RT offset (β_0_), and a stronger influence of unexpected uncertainty (β_4_). Further model-based simulations indicated that this parameter profile reflects reduced precision-weighting of prediction errors in older adults, consistent with diminished stimulus-driven updating and a greater reliance on prior expectations. These findings demonstrate that cognitive control mechanisms contribute to postural and stepping behaviour and that age-related changes in these processes result in reduced adaptability that may increase fall risk in older age.

In model-free analysis on empirical data, consistent age-related effects were observed in all four force latency measures (APA onset, maximum APA, lift-off, and touchdown times) but were absent in movement time. This demonstrates that the observed differences arise at an early, anticipatory stage of step initiation rather than reflecting alterations in movement execution.

These findings on delayed response times were consistent with previous studies looking at changes in postural control and gait initiation with age ^16,36–40^. We also found an increase in APA onset time at mostly-no-go compared to mostly-go conditions, which was again consistent with a previous gait initiation study where go/no-go expectancy was experimentally manipulated ^17^. In addition, older adults showed reduced trial history dependence while anticipating a stepping response, and this age-related difference prominently seen in mostly-go condition was highlighted by a steeper decrease in APA onset time with trial history in younger adults. Such trial history dependent changes were consistent with accounts on usage of serial dependency in perceptual tasks with repetitive stimuli ^18,41^ and were in line with previous findings from neurocognitive studies on effects of serial dependency in ageing ^42,43^. These empirical RT results on age-related slowing and differences in trial history dependence suggest that changes in anticipatory processes contributed to altered gait initiation, reflecting the involvement of early brain processes and cognitive control.

Computational modelling allowed us to disentangle these anticipatory cognitive control processes in gait initiation using three HGF model parameters: the learning rate (ω), RT offset (β_0_), and sensitivity to unexpected uncertainty (β_4_). Age-related differences in ω captured reduced statistical learning in older adults. In previous studies that applied HGF to behavioural data ^44,45^, reduction in ω suggested a diminished updating of higher-order perceptual beliefs. By linking age-related differences in ω to statistical learning, our results suggest that younger adults dynamically updated predictions by leveraging probabilistic trial dependencies, prioritizing accumulated sensory evidence under environmental uncertainty. In the response model, the differences in β_0_ reveals a general age-related increase in RT, while β_4_ captured slower RT responses in older adults to unexpected alterations in the stimulus sequence. In a previous study employing the same model, β_4_ was linked to post-error slowing ^45^. In our data, this age-related increase in β_4_ indicated a diminished tendency for pre-error speeding to unexpected changes in go/no-go sequences, which may reflect strategic slowing and cautious responses to competing go/no-go stimuli ^46^ – consistent with uncertainty underestimation that weakens adaptive learning from probabilistic feedback ^32^.

Simulations of perceptual hidden states showed larger precision weighting (ψ) on prediction errors (ε) in younger adults, and this pattern was stronger from the first simulated trial up until the switch in probability of go/no-go trials. Based on the predictive coding framework ^23,47^, these simulated patterns of ψ and ε reveal distinct adaptive strategies in ageing. Younger adults show a transient rise in precision weights at the start of the stepping task, followed by a decrease in precision weights closer to the switch of conditions. This pattern of precision weighting in younger adults reflects a more adaptive usage of incoming stream of statistical information from the environment to revise anticipatory predictions. In contrast, the flatter precision profiles in older adults point to reduced sensitivity to environmental changes, showing a stronger reliance on prior beliefs when initiating and timing stepping responses. Together, these findings reveal an age-related shift from rapid online adjustments to changing sensory inputs toward belief updating guided by greater reliance on lifespan-accumulated priors ^27,30–32^. Under the free energy principle, this age-related shift enhances cortical efficiency and perceptual stability ^48^.

Reduced belief updating suggests that older adults tend to become less exploratory, showing reduced cognitive flexibility and more rigid posterior expectations ^27^. Such processes in postural control and gait may end up being maladaptive as they can reduce the efficiency with which prediction errors are integrated to refine the individual’s forward model. This could lead to cumulative prediction errors that degrade the precision of anticipatory adjustments, thereby forcing older adults to rely more heavily on reactive control to maintain stability and balance.

Learning rate and unexpected uncertainty were inferred from trial-wise dynamics with only a single change-point and fewer trials in older adults (160 vs 240), which may impair parameter identifiability^49^. Given the limited volatility associated with a single-change point in outcome probabilities, we therefore used a reduced two-level HGF. Multiple change-point switches could be incorporated to better assess volatility’s impact on perceptual learning rates^50^ in gait initiation, enabling examination of belief updating driven by volatility estimates in a three-level HGF^29^. For example, previous neurocognitive studies have manipulated environmental volatility by altering stimulus variance in motion tasks ^51^ or by changing reward probabilities ^21^ to examine its role in dynamically updating learning rates and salience on statistical information when uncertainty is high. Dynamically changing environmental volatility would better explore the role of volatility in dynamically updating precision weights on prediction errors, offering deeper insights into neural mechanisms contributing to adaptive exploratory learning such as the ACC ^21,52^, which is particularly relevant given age-related cognitive declines in decision-making have previously been associated with ACC ^53^. Furthermore, including additional tests such as N-back for proactive interference ^54^, Stroop/Flanker for cognitive conflict resolution and error slowing ^55^, Wisconsin Card Sorting Test (WCST) for set-shifting ^56^, or Balloon Analogue Risk Task (BART) for risk under uncertainty ^57^ would offer a more nuanced understanding of cognitive control processes and age-related cognitive flexibility. Including these in future studies would enable to look at correlations across subjects between model parameters and cognitive control metrics, forging direct links between model parameters and behavioural measures of cognitive flexibility.

We show reduced precision-weighting of prediction errors in older adults through computational modelling of behavioural data. Future studies could complement this approach by adding neurophysiological measures such as mobile EEG or fNRIS^58^. Computational modelling on APA onsets provided indirect mechanistic insights into age-related anticipatory control by inferring latent variables, while neuroimaging may assess precision-weighting of prediction errors more directly. For example, an electrophysiological study looking at cognitive control in gait has shown how modulations in beta-band oscillatory networks across parietal cortex and PFC play a crucial role in top-down signalling for flexible adjustment of behaviour in anticipation of a change in auditory pacing of cues^59^. For temporal predictions observed in anticipatory processes, an elevation in beta synchronization indicates priority of the brain to maintain stability by actively maintaining probabilistic priors accumulated over the long-term whereas a desynchronization signals disruption of current sensorimotor or cognitive set by unexpected novel events ^60^. Consistent with predictive coding accounts of motor control, sensorimotor beta activity has been shown to scale with the precision-weighting of prediction errors during action, linking uncertainty-driven belief updating to exploratory learning and movement execution across visuomotor tasks ^61,62^. Future studies could integrate mobile EEG with go/no-go stepping paradigms to determine whether sensorimotor beta dynamics index age-related changes in the precision-weighting of prediction errors during step initiation.

In summary, our findings reveal age-related differences in anticipatory postural control by separable cognitive control processes. Older adults showed slower postural onset responses, reduced statistical learning, and diminished pre-error speeding, relying more on empirical priors and less on serial dependencies. Younger adults, by contrast, dynamically tuned responses via proactive control using environmental statistics. These findings align with predictive coding accounts that favour stable top-down predictions over flexible sensory integration with age, highlight how hierarchical learning shifts modulate proactive-reactive control in postural anticipation. Future mobile brain-body imaging studies should clarify underlying oscillatory dynamics involved in cognitive control of adaptive postural control in ageing.

## 4. Methods

### 4.1. Participants

In this study, 24 younger adults aged 20 to 40 years and 25 older adults aged 65 to 85 years with a BMI ranging between 18 to 25 kg/m² were recruited. The younger group consisted of 19 female and 5 male participants, and the older group consisted of 16 female and 9 male participants.

Participants with presence of pain in any of their body parts, musculoskeletal conditions affecting sit-to-stand performance, and neurological disorders were excluded.

Given the final sample size, number of measurements, an alpha level of 0.05, and a desired power of 0.80, a posteriori sensitivity analysis ^63^ indicated that the smallest effect size (Cohen’s f) detectable with the sample was 0.41, which corresponds to medium to large effect.

Participants for this study were recruited either at Maastricht University, or they lived within the region of Maastricht or the province of Limburg, the Netherlands. The study was approved by the Ethics Review Committee Psychology and Neuroscience at Maastricht University (ERCPN-OZL_231_155_12_2020), and all participants gave written informed consent before participation.

### 4.2. Experimental Procedure

At the beginning of the study, body measures such as height, weight, and BMI were recorded from the participants in both age groups. Participants were asked to indicate which leg was dominant (left/right). We used the Berg Balance Scale (BBS) to screen older participants for fall risk ^64^. A score of 45 points or more indicates that the participant can walk safely and independently, and we used this score as a cutoff for study participation. To assess functional mobility ^65^, all participants performed a Timed-Up and Go test (TUGT).

Participants performed a choice stepping task using a go/no-go paradigm, implemented on a computerized stepping mat containing eight panels: two centre panels, two forward panels, two backward panels, and one panel each on the left and right side respectively ^66^. Participants were standing on the centre panels and stepped forward dependent on the stimuli that were presented on a computer monitor, placed 2 meters in front of the stepping mat at a height of 1.5m (Fig. 1A).

For the go/no-go paradigm, each trial consisted of a baseline interval (1500 ms) during which a fixation cue was presented, followed by a spatial cue (500 ms) that indicated with which foot the step should be made. Following a variable inter-cue interval (1000-2000 ms), a go or no-go cue (2000 ms) was presented (Fig. 1B). Following a go cue, participants had to step onto the forward panel, and following a no-go cue, they had to remain standing on the centre panels. The trial ended with a recovery period of 1000 ms for a no-go trial or when both feet were positioned back to the centre panels in the case of a go trial. Participants were asked to maintain their gaze onto the fixation cross at all times.

We varied the odds ratio between go and no-go trials to experimentally manipulate the anticipation of movement. The ratio of go and no-go trials was set to 3:1 (75%) in the mostly-go condition and 1:3 (25%) in the mostly-no-go condition. The 3:1 expectancy ratio was selected to maximize response conflicts by increasing false alarms ^67^. Both conditions were presented in separate blocks. In younger participants, the experiment was divided into two blocks of 120 trials each (240 trials in total). To reduce the overall load, older participants performed two blocks of 80 trials each (160 trials in total). Within each block the order of trials was randomized for each participant separately with equal allocation of left and right go and no-go trials in each block, while the order of the blocks was counterbalanced between participants. There was a short break after every 10^th^ trial, and participants were instructed to realign their foot positions to the centre panels of the stepping mat. After each quarter of the total number of trials, participants were given a longer break.

A custom-made script was written using Presentation® software (Version 23.0, Neurobehavioral Systems, Inc., Berkeley, CA, www.neurobs.com) for visual stimulus presentation and recording responses from the stepping mat.

### 4.3. Data Acquisition

During the study, three-dimensional ground reaction forces (GRF) were measured using two force plates of dimensions 60 × 40cm (AMTI OR6-Series, Watertown, USA; 1kHz) at a sampling rate of 1 kHz. These were placed underneath the stepping mat so that one force plate was below the left centre and forward panels and the other under the right centre and forward panels. The data were stored in a coordinate 3D format (C3D) using Nexus software (Vicon Motion Systems, Oxford, UK).

Only the centre panels and forward panels of the stepping mat were used. The centre panels had a diameter of 20cm, and the forward panels had a dimension of 19.5 × 20cm. The centres of both panels were separated by a length of 28.75cm. Sensors placed inside each of the panels of the computerized stepping mat recorded step responses such as foot lift-off and touchdown.

The step response data from the stepping mat were recorded by the stimulus presentation software together with stimulus presentation timings at 1ms precision and stored as unique markers in the log file.

### 4.4. Data Analysis

All trials were visually inspected to reject duplicates and missing event markers. Lift-off time was quantified as the difference between centre panel release time and go cue presentation time, and touchdown time as the difference between forward panel touchdown time and the go cue presentation time. The movement time was quantified as the difference between forward panel touchdown time and centre panel release time.

Vertical force data of the stepping leg and stance leg were extracted from the two force plates. From the vertical GRF, anticipatory postural adjustments (APA) were determined before the foot lift-off. APA is a form of control strategy which activates postural muscles prior to a stepping response to maintain equilibrium ^68^. Maximum APA was defined as the postural event when the magnitude of vertical GRF of the stance leg between go cue and lift-off reached a minimum value ^69,70^. Anticipatory postural onset (APA onset) was defined as the postural event between the go cue and maximum APA when the derivative of the force differences between the stance leg and the stepping leg reached zero. Both the latency and the normalized vertical force were assessed for the APA onset and maximum APA respectively (see Section S1 and S2 for more details).

Not all trials showed an APA. To identify trials with an APA, a threshold was set to determine the presence of an APA ^15^ and to determine the latency of maximum APA and APA onset. First, the derivative of the low-pass filtered (50 Hz) vertical force difference between stepping leg and stance leg was computed. The threshold was defined as five times the standard deviation of the derivative, calculated over the time window from 100ms before to 25ms after the go cue. When the derivative exceeded this threshold, the maximum APA was identified as the peak vertical force of the stepping leg after reaching the threshold and before foot lift-off, based on the raw force signals. APA onset was defined as the point at which the derivative of the vertical force difference first crossed zero before reaching the threshold (Fig S1.2A). All trials with APA were reviewed, with special attention to those at the tail ends of the force latency histograms (Fig S1.2B). From this visualization, trials in which participants shifted the weight distribution between legs before the go cue were excluded.

### 4.5. Statistical Analysis

RT data (APA onset, max APA, lift-off and touch-down latencies) and normalized vertical GRF of stepping leg at Max APA were statistically analysed using a Linear Mixed Model (LMM). In this model, the fixed effects variables were condition (mostly-go/mostly-no-go) and group (Young/Old). The random effects group factors were set to participant ID. Post-hoc analysis was done by specifying contrasts within the estimated marginal means (EMM) function and Bonferroni correction was used to adjust p-values for multiple comparisons.

To assess serial dependencies in RTs, trial history was quantified as the number of consecutive go or consecutive no-go trials: if a go trial happened after 3 consecutive no-go trials it was labelled as ‘-3’, whereas if a go trial was the 4^th^ consecutive go-trial it was labelled at ‘4’. Here, we applied a LMM on APA onset time with fixed effect variables trial history (continuous) and group (Young/Old).

Statistical significance threshold was set to 0.05. Cohen’s d effect sizes and 95% confidence intervals were estimated for t-tests, and partial eta squared effect sizes, and 90% confidence intervals were estimated for the LMMs. Statistical analyses were done in JASP (https://jasp-stats.org/) and R.

### 4.6. Computational Modelling

A Bayesian modelling approach was used to estimate hidden parameters associated with the serial dependencies in RT data. To look more closely at how participants update their internal model based on trial history, it is necessary to understand how uncertainty of sensory information in the environment, i.e., the probability of sequential go/no-go cues, affect the participant’s perceptual beliefs and how these beliefs change with behavioural responses on a trial-to-trial basis. To this end, we used a Hierarchical Gaussian Filter (HGF). The HGF is an empirically validated hierarchical decision model designed to test belief updates under different sources of uncertainty within an adaptive environment using mean-field approximations and full Bayesian solutions ^28,29^.

The HGF consists of a perceptual and a response model. The perceptual model tracks trial-wise learning during a task, and the response model translates the perceptual beliefs into task responses in a trial-wise manner (see Fig. 2A). In this study, we chose the enhanced HGF for binary outcomes as the perceptual model, and a calibrated linear log RT model as the response model ^45^. Since the experiment had only a single change-point in outcome probabilities across trials and didn’t manipulate volatility, we chose a two-level enhanced HGF. More details about the model equations, calibration, and prior parameters are provided in Section S3 of the Supplementary Information.

To quantify the extent to which statistical learning supported the online perceptual anticipation of sequential go/no-go events within each participant, a learning rate ω was extracted from the perceptual model. This parameter captures the tonic degree to which existing prior beliefs are updated by new information from the environment, in proportion to the relative precision of new evidence compared to that of the current beliefs (see Equation S3.3)^29^. Typically, ω increases when environmental uncertainty rises, causing participants to place greater weight on newly accumulated evidence to update their internal beliefs. Therefore, this term captures the step size of change in probabilities across stimulus transitions and the rate at which posterior expectations are updated.

The response model contained 6 free parameters (see Supplementary Information, Section S3). Of these parameters, parameter β_4_ is of special interest as it captures the weight of unexpected uncertainty (phasic volatility) on RTs. Since log RT is proportional to estimates of unexpected uncertainty weighted by β_4_, higher β_4_ translates to slower RTs.

We fitted the resulting model to individual participants’ trial-by-trial APA onset times as the participant responses and the go/no-go cues as the binary stimulus inputs. Model fitting was performed using the Broyden-Fletcher-Goldfarb-Shanno algorithm as implemented in the HGF Toolbox. By fitting the model, parameters including learning rate ω and β parameters were estimated for each participant.

To test whether the estimated model parameters differ between groups, an independent samples t-test was performed for each parameter separately and Benjamini-Hochberg correction was used to adjust p-value for multiple comparisons. Posterior predictive checks were conducted across the parameter space to examine how variations in model parameters related to the observed empirical results of APA onset times and trial history. The perceptual model of the simulated HGF was then used to generate the latent variables of precision weights (ψ) on prediction errors and precision-weighted prediction errors (ε), based on the mean parameter values (ω and β_4_) extracted from participants.

## Data availability

Data and code for the analyses presented in this paper are available at https://github.com/thebince/Reduced_BeliefUpdating_OlderAdults_StepInitiation

## Acknowledgments

We would like to thank Li-Juan Jie, Chloé Calis, Helge Monie, Floor Kaalberg, Allanah Smith, Pien Alferink, Jeannette Boschma and Penelope Edwins for their assistance with data acquisition, and Johan Gielissen and Paul Willems for technical support.

## Author contributions

T.W.B., S.A.K., K.M., and M.K. were involved in design of the study. M.K. collected the data. A.B.J. was involved in data curation. A.B.J., R.A., and T.W.B. were involved in data analysis and investigation. A.B.J. wrote the original draft of the manuscript. A.B.J., T.W.B., K.M., S.A.K., R.A., and M.K. were involved in reviewing and editing of the manuscript.

## Supplementary Information

**Table S1.**
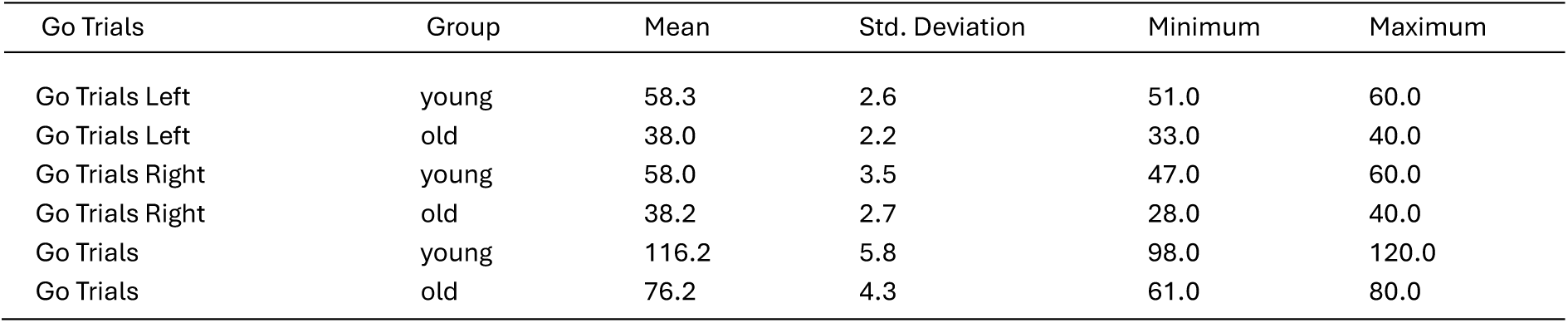
Descriptive statistics of the number of Go trials included in the analysis.

### S1. Normalized Vertical Force

Vertical ground reaction force of stepping leg and stance leg were normalized to body weight (BW). BW can be estimated as the force of both legs averaged over time.

**Fig. S1.1.**
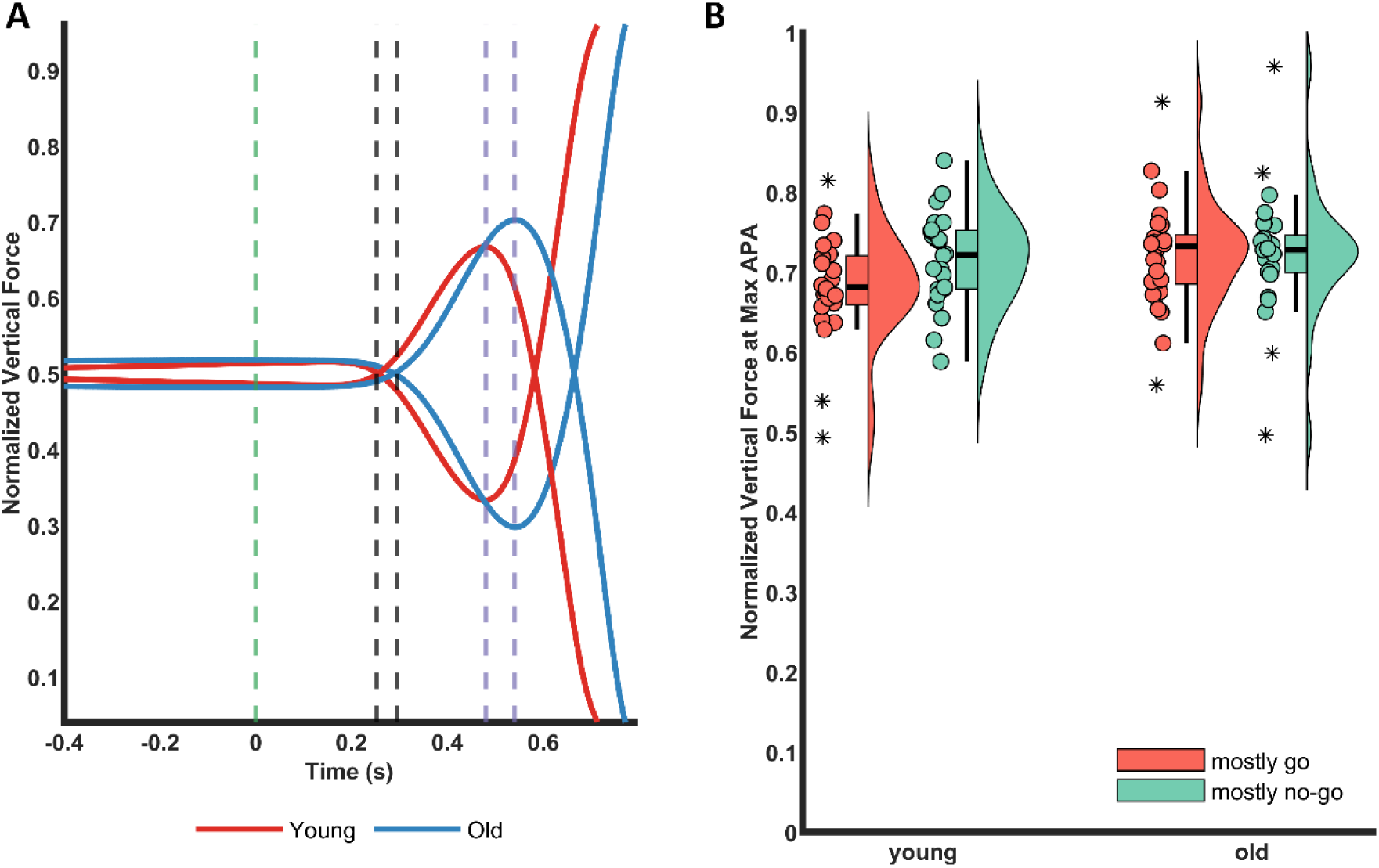
Force Averages. **(A)** Average vertical forces of young and old normalized to body weight. The green line indicates go cue, black lines indicate APA onsets, and purple lines indicate maximum APAs. On average, older people show a higher maximum APA than younger people. **(B)** Raincloud plots of normalized vertical force of stepping leg at maximum APA as a function of condition (mostly go / mostly no-go) and groups (young/old).

**Fig. S1.2.**
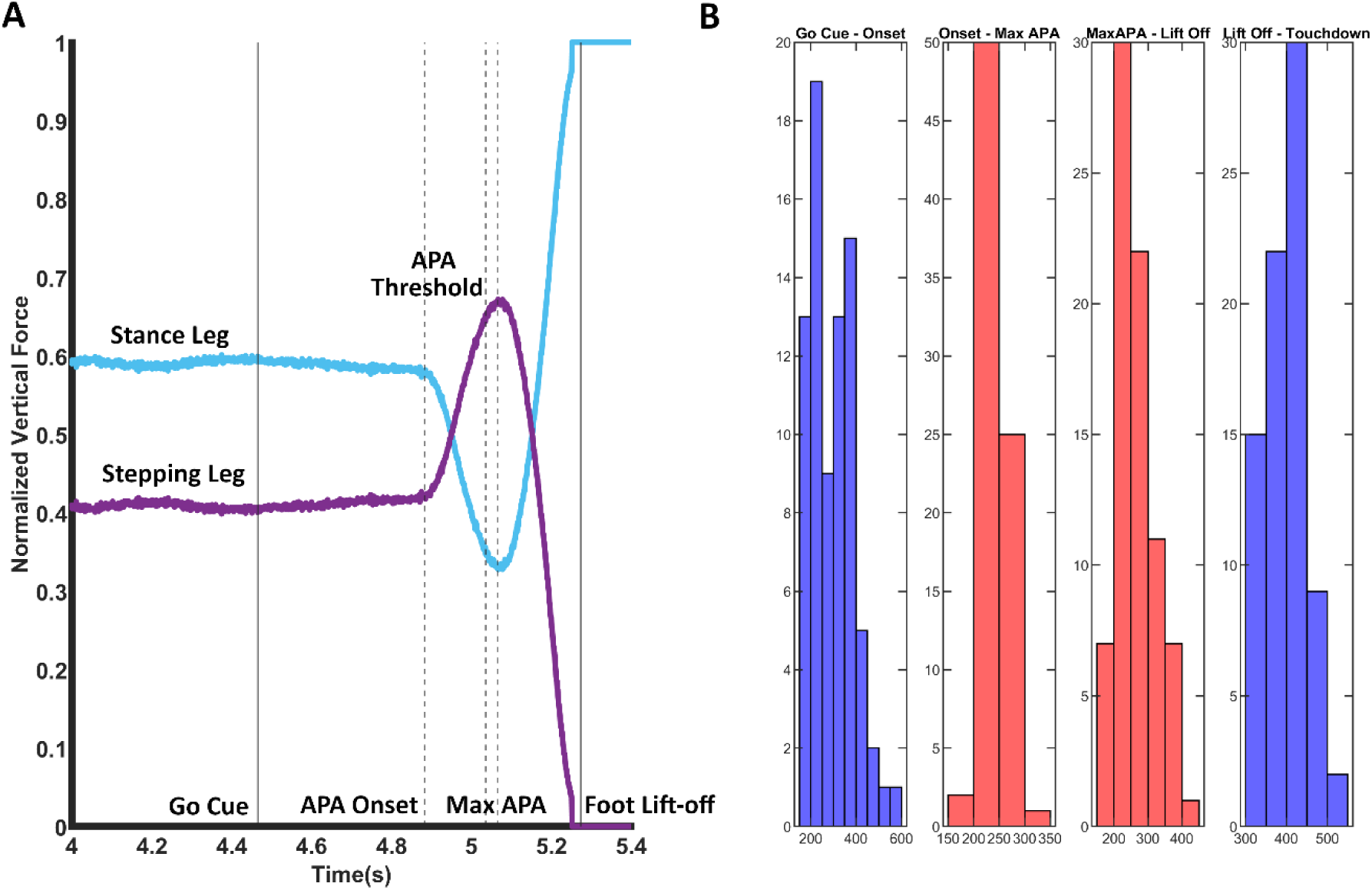
Maximum APA Detection. **(A)** Normalized vertical forces of stepping leg and stance leg in a go trial of an example participant. Here, the APA threshold was used to identify the maximum APA and APA onset. Maximum APA was identified as peak vertical force of stepping leg after the threshold and before foot lift-off. APA onset was identified as the first zero crossing point of the derivative before threshold **(B)** Histogram of a participant showing the distribution of latencies from go cue to APA onset, APA onset to maximum APA, maximum APA to foot lift-off, and foot lift-off to touchdown. The latencies of force trials present in the tail ends of the distribution were visualized to see if they were outliers or not.

**Table S1.1.**
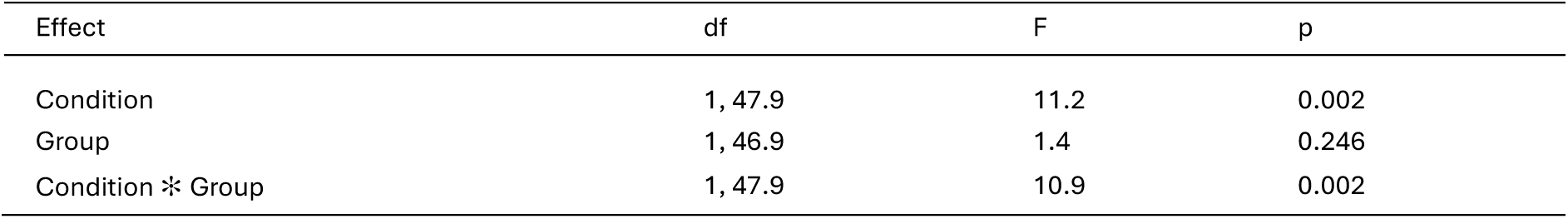
ANOVA Summary of Linear Mixed Model on Normalized Vertical Force of Stepping Leg at Maximum APA.

**Table S1.2.**
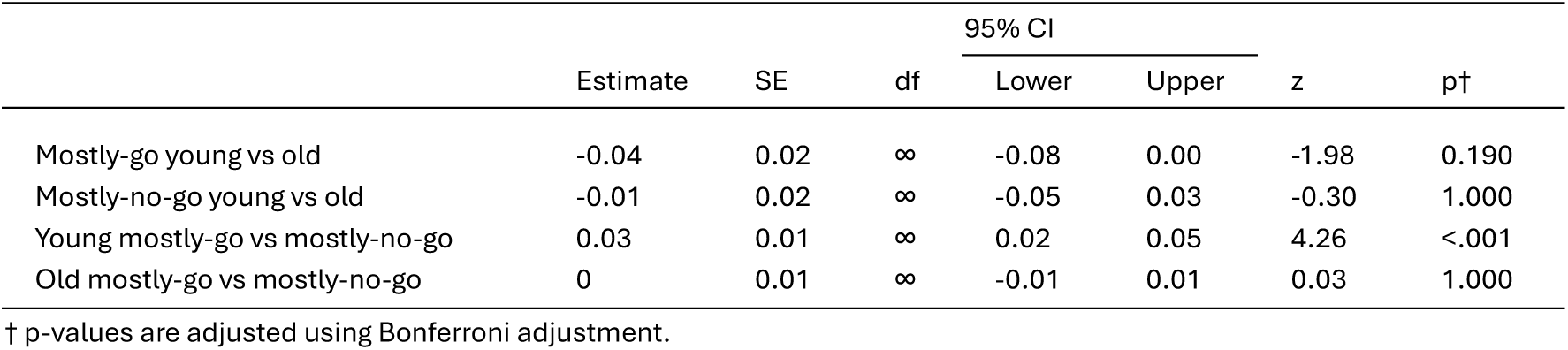
Contrasts of Linear Mixed Model on Normalized Vertical Force of Stepping Leg at Maximum APA.

### S2. Force Latencies and Trial History

**Fig. S2.1.**
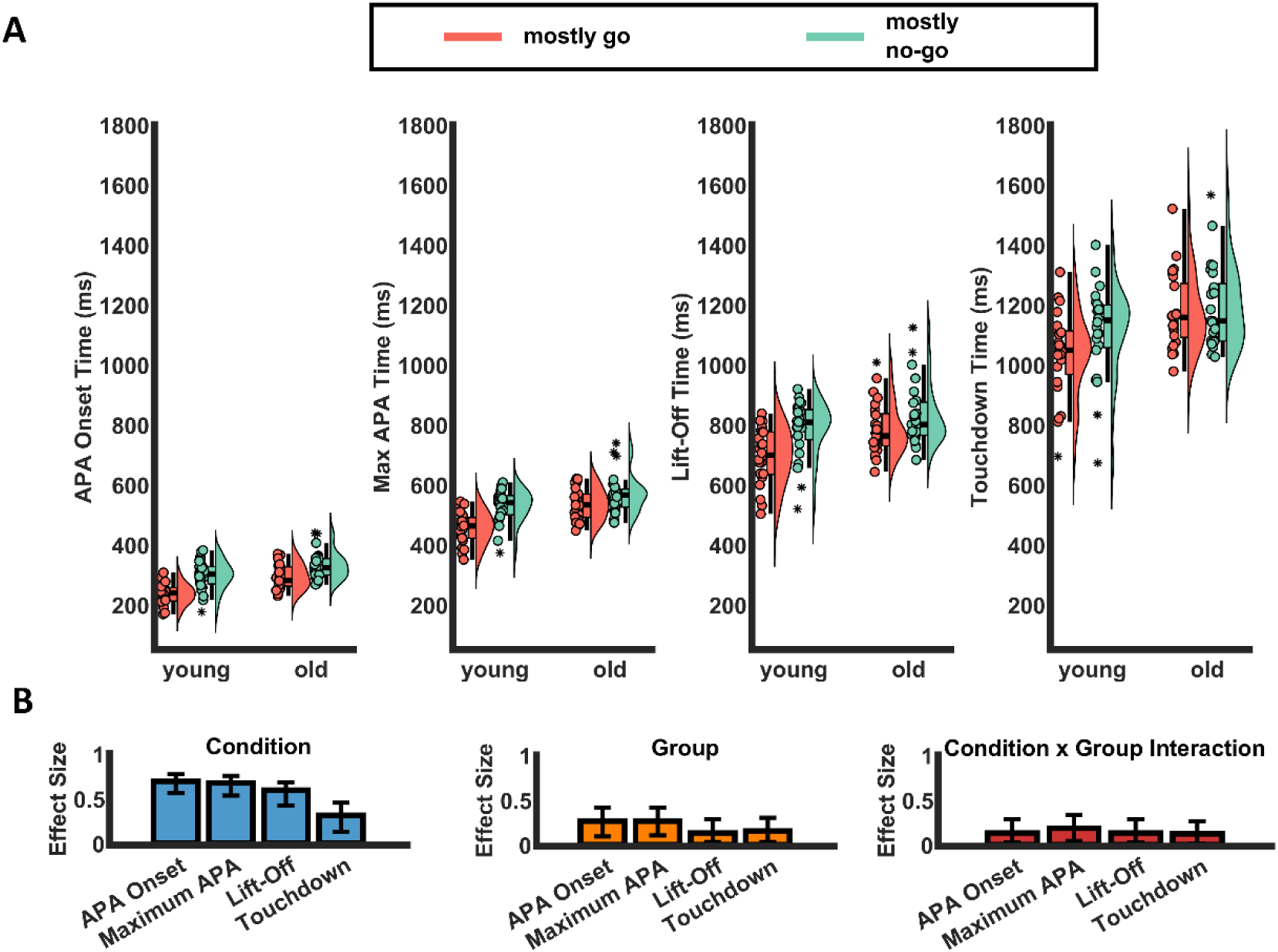
Reaction Time Descriptive Plots. **(A)** Raincloud plot of APA onset time, Maximum APA time, Lift-off time and Touchdown time of young and old participants. The scatter plot indicates individual time points, colours indicate conditions, and the whiskers indicate outliers **(B)** Bar plot of the estimated effect sizes (η² partial) of condition, group, and interaction for each of the four time points. Error bars indicate the 90% confidence interval.

**Table S2.1.**
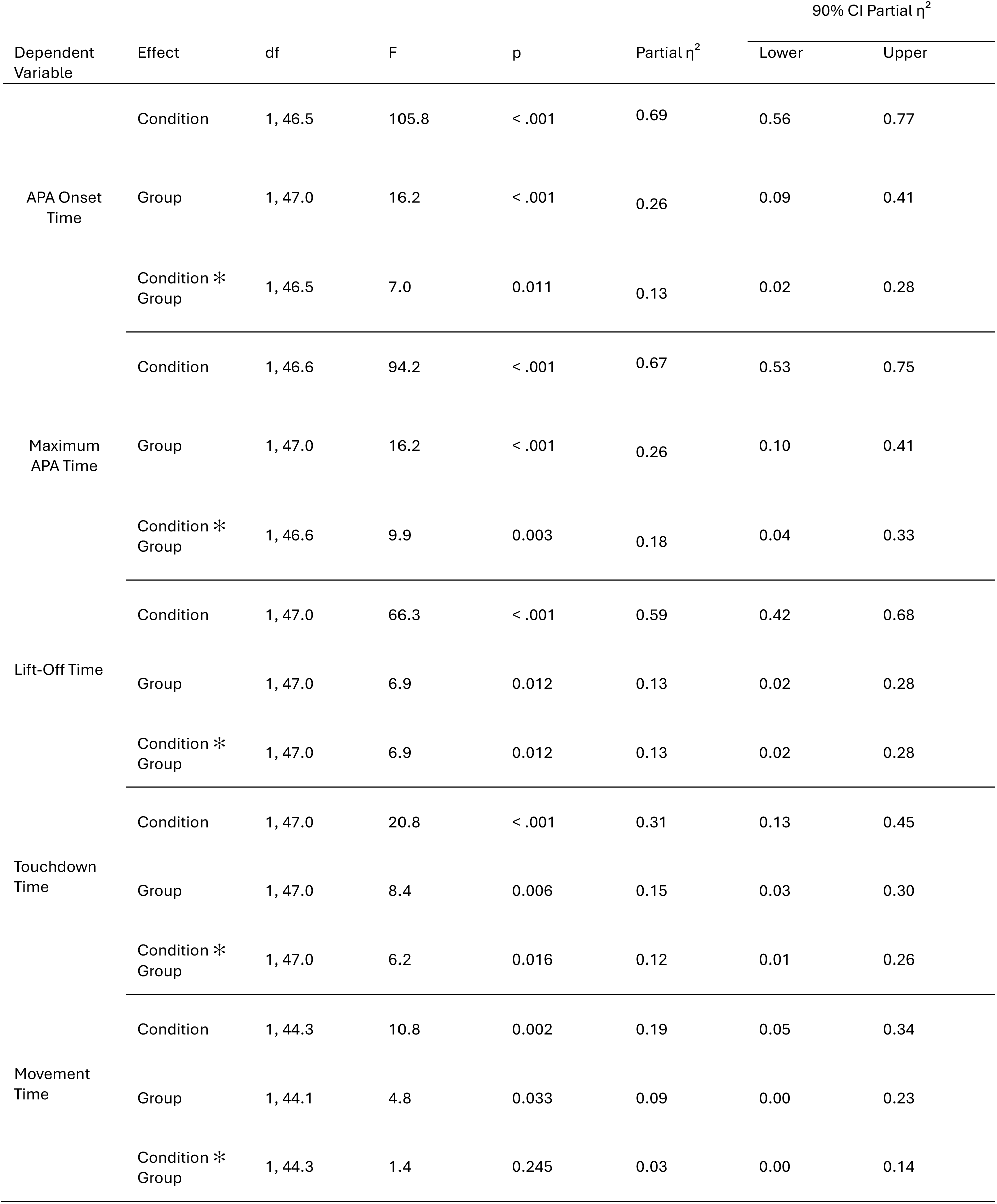
ANOVA Summary of Linear Mixed Model of APA Onset Time, Maximum APA Time, Lift-Off Time, Touchdown Time and Movement Time.

**Table S2.2.**
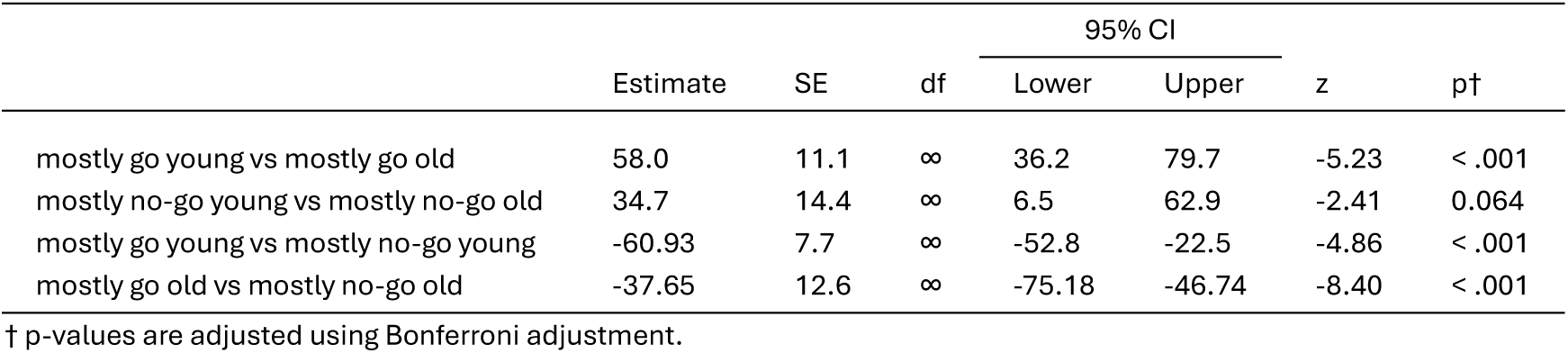
Contrasts of Linear Mixed Model on APA onset time.

**Fig. S2.2.**
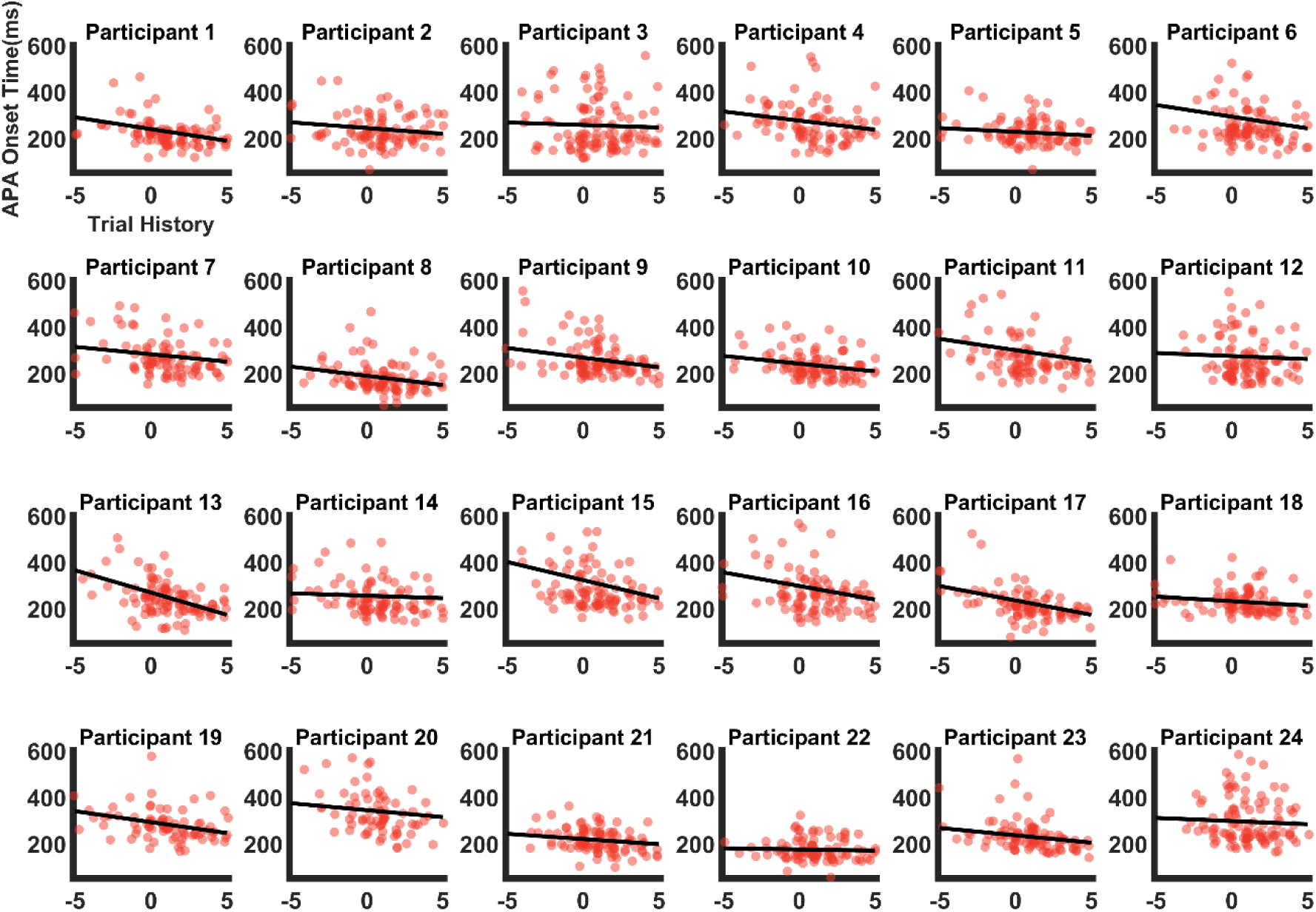
Scatter plots of the APA onset times of individual trials a function of trial history of the young participants

**Fig. S2.3.**
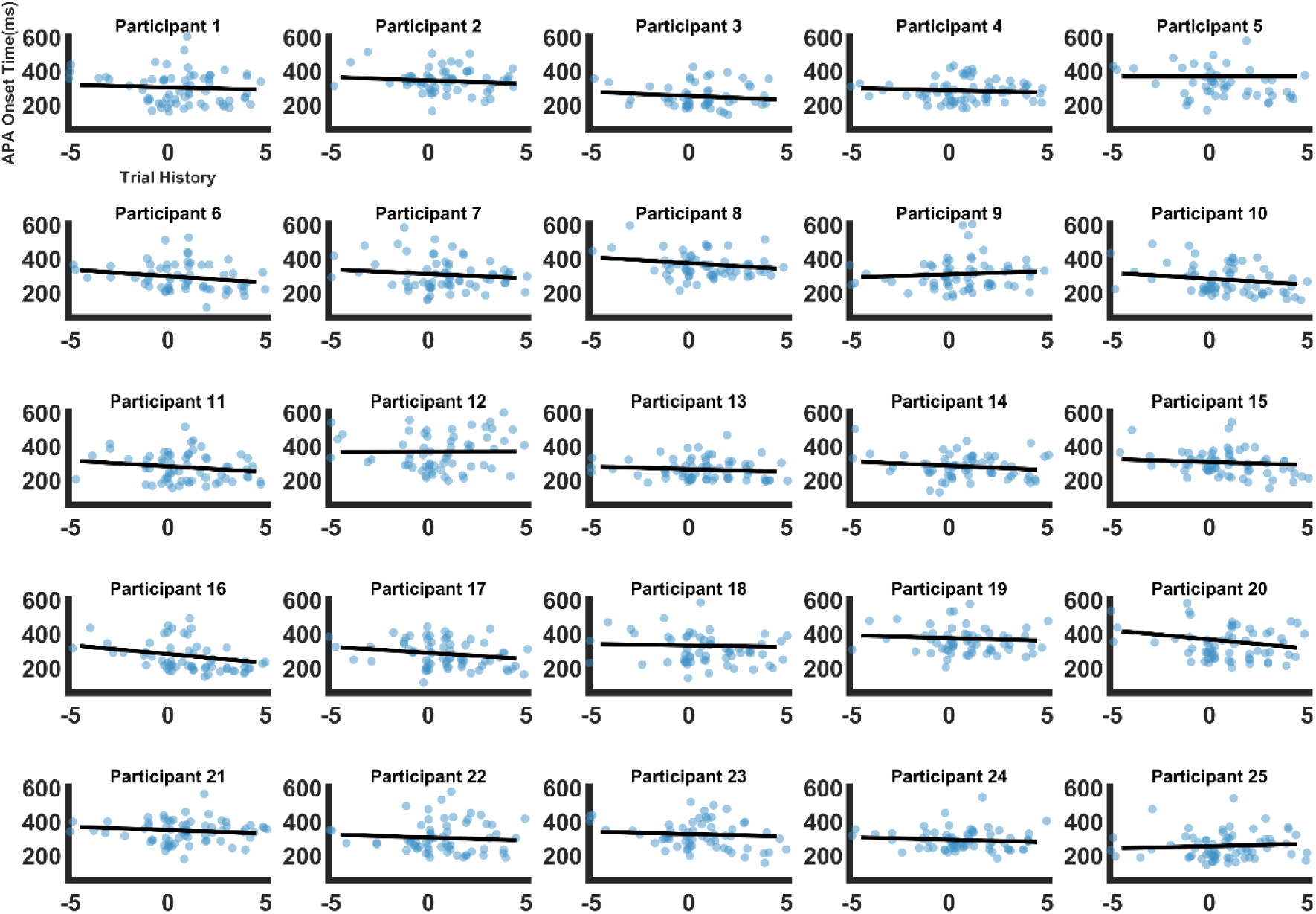
Scatter plots of the APA onset times of individual trials a function of trial history of the old participants

**Table S2.3.**
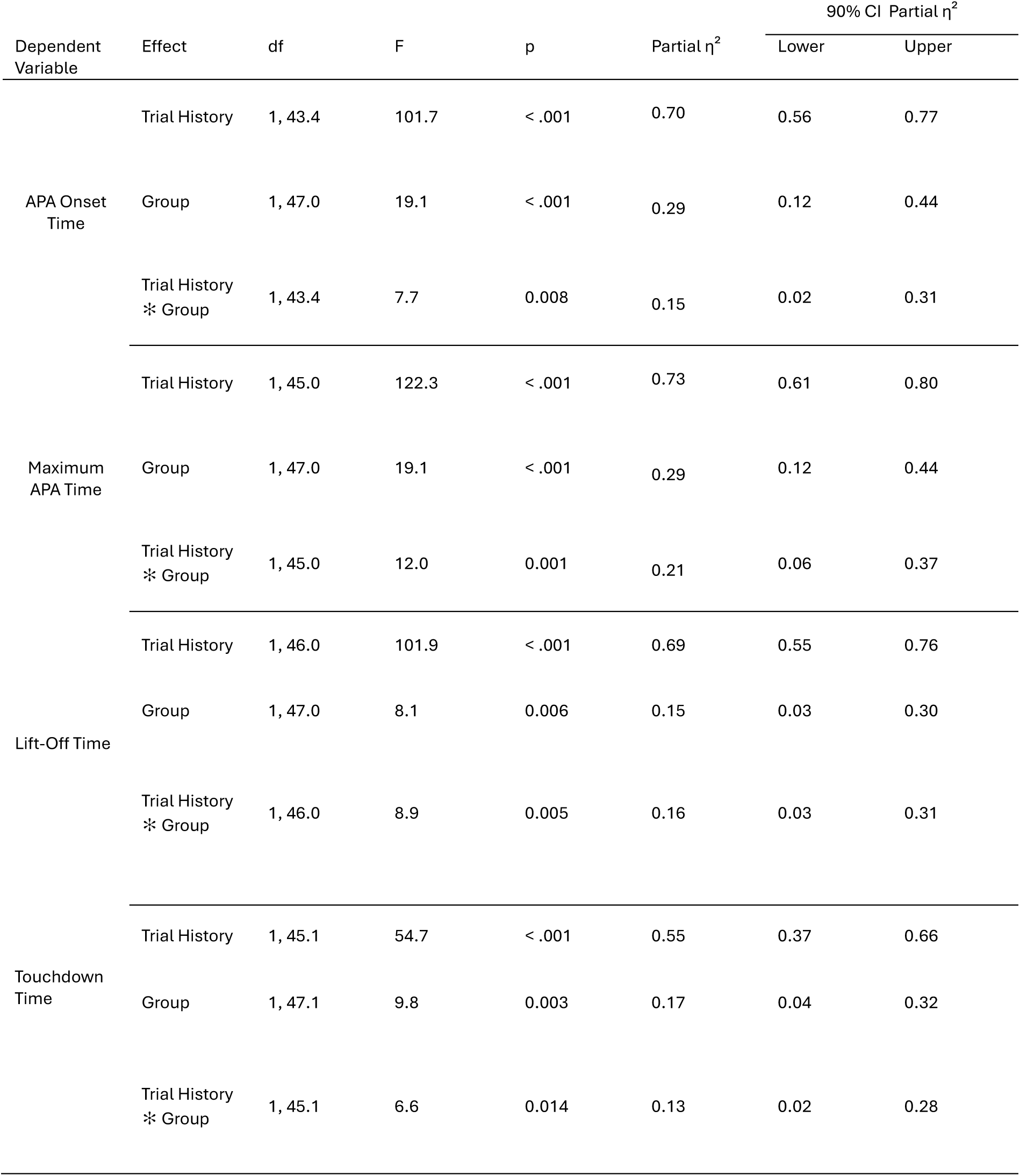
ANOVA Summary of Linear Mixed Model on Trial History on APA Onset Time, Maximum APA Time, Lift-Off Time, and Touchdown Time.

### s3. Hierarchical Gaussian Filter

The response model used in our study is a log-reaction time model (Log (RT)). Log (RT) is described as a function of six gaussian parameters: β_0_, β_1_, β_2_, β_3_, β_4_, and ζ. These coefficients of the response model describe how trial-wise beliefs are translated into log-reaction times. The parameters β_0-4_ capture the weights of surprise of the stimulus outcome (Go/No-Go), certainty of the outcome, probability of informational uncertainty, environmental uncertainty, and Gaussian decision noise on the predicted log (RT). The default equation of the Log RT model is given as follows:

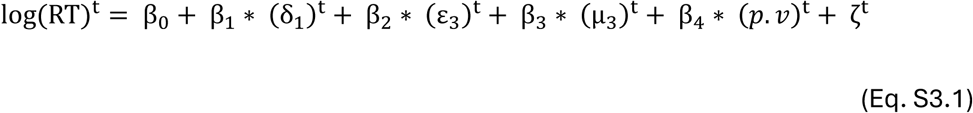

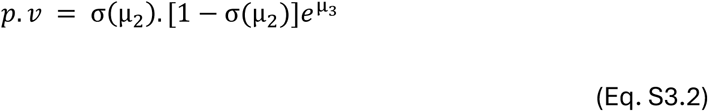

In Eq. S3.1, δ_1_ is surprise, ε_3_is Bernoulli variance, μ_3_ is inferential variance, and 𝑝. 𝑣 is phasic volatility (Eq. S3.2). Because the task structure and RT distribution deviated from canonical paradigms previously fitted with HGF, we first calibrated the model by performing a prior predictive check of the HGF linear response model for log RTs. Specifically, we simulated RTs across a range of response model parameters while keeping the perceptual model fixed to Bayes-optimal values. We compared the simulated RT distributions to the empirical RT density averaged across participants, adjusting the response model’s prior means until the simulated distributions qualitatively reproduced the empirical RT distribution. These tuned priors were then used as group-level priors in subsequent model fitting and between-group comparisons. The resulting prior parameter values were as follows: β_0_ (μ = log (200); σ = 4), β_1_ (μ = 0; σ = 4), β_2_ (μ = −2; σ = 4), β_3_ (μ = 2; σ = 4), β_4_ (μ = 2; σ = 4), and ζ (μ = log (log (20)); σ = 4).

The tonic learning rate ω is a function of outcome uncertainty, informational uncertainty, and environmental uncertainty. The equation of learning rate ω is given as follows:

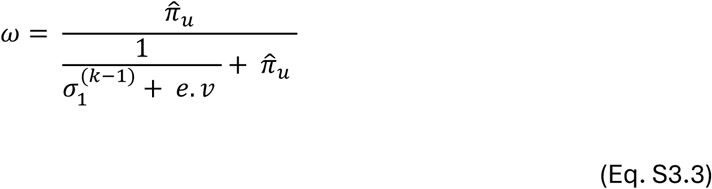

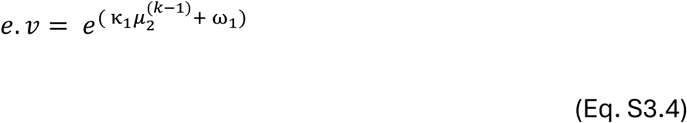

In Eq. S3.3, 𝜋^_𝑢_ is the outcome uncertainty, 𝜎 is the informational uncertainty, and 𝑒. 𝑣 (Eq. S3.4) is the environmental uncertainty.

**Table S3.1.**
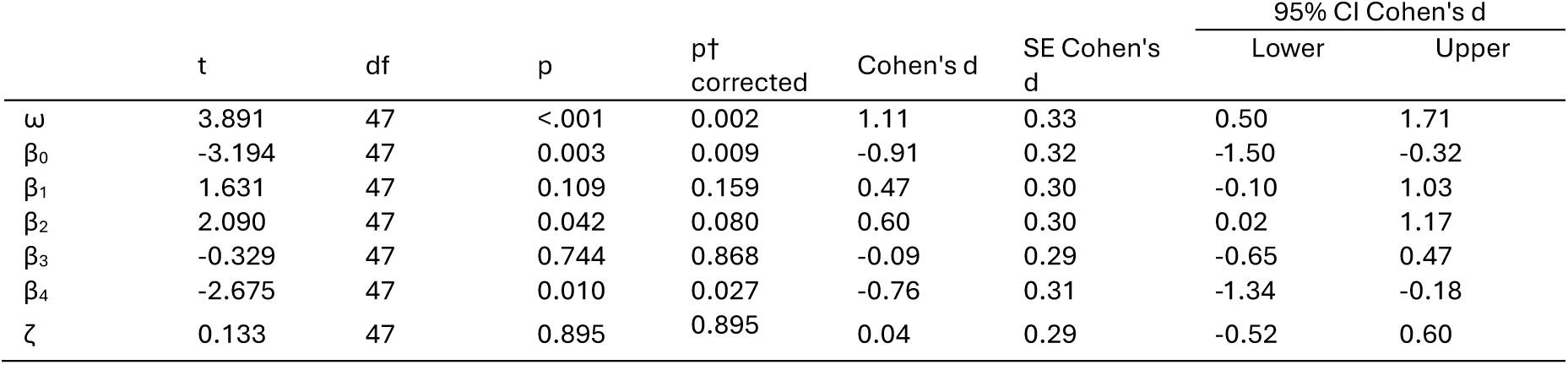
Independent samples t-test of perceptual and response model parameters.

**Table S3.2.**
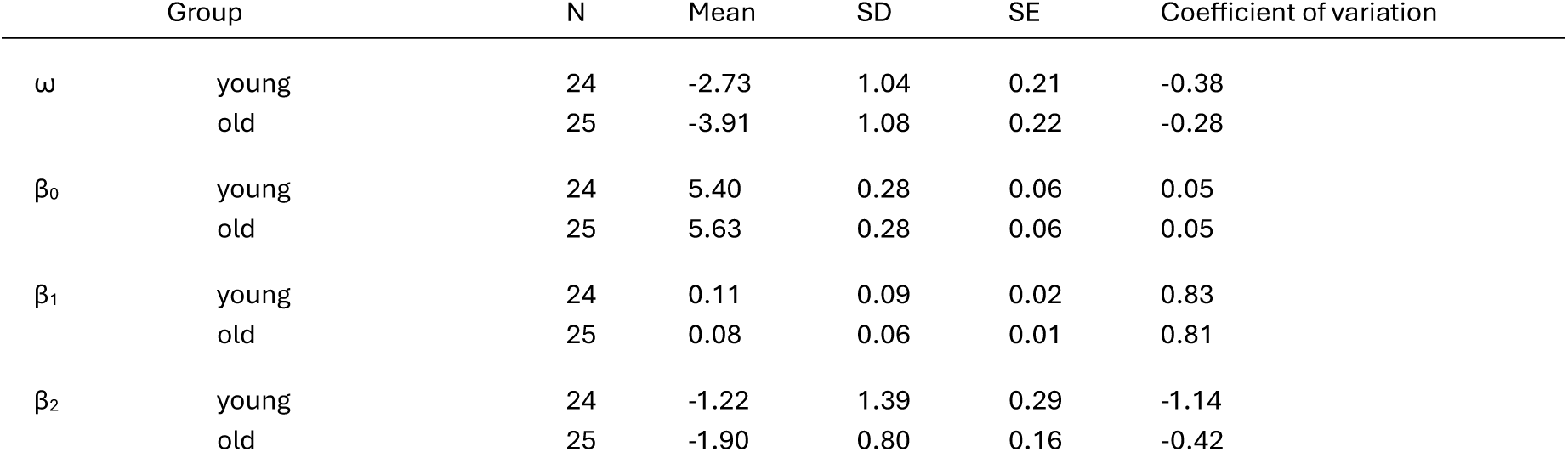
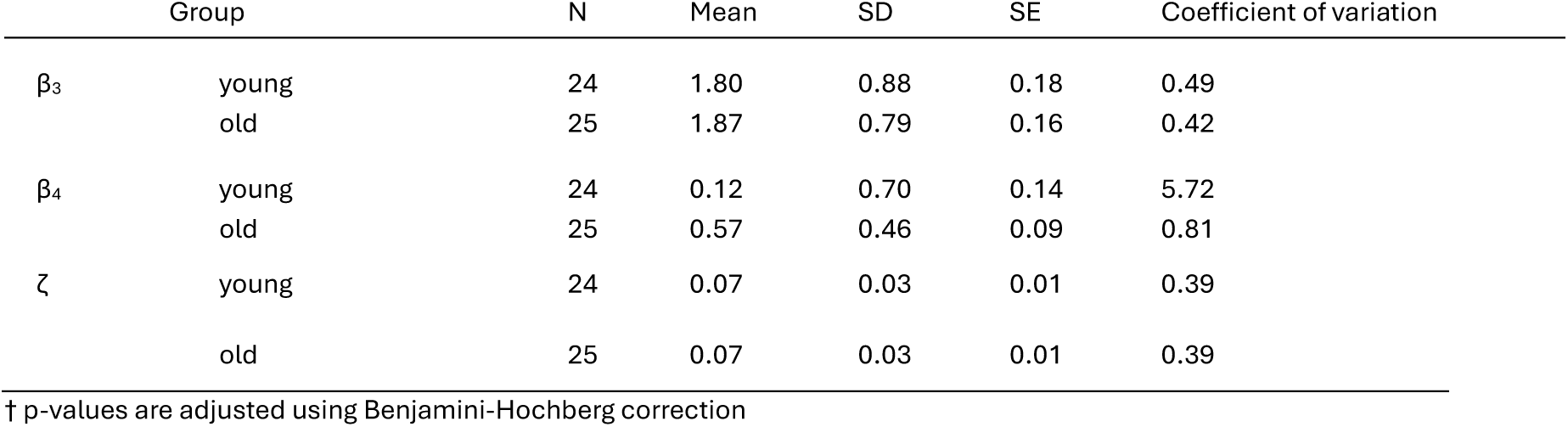
Descriptive Statistics of perceptual and response model parameters.

**Fig. S3.1.**
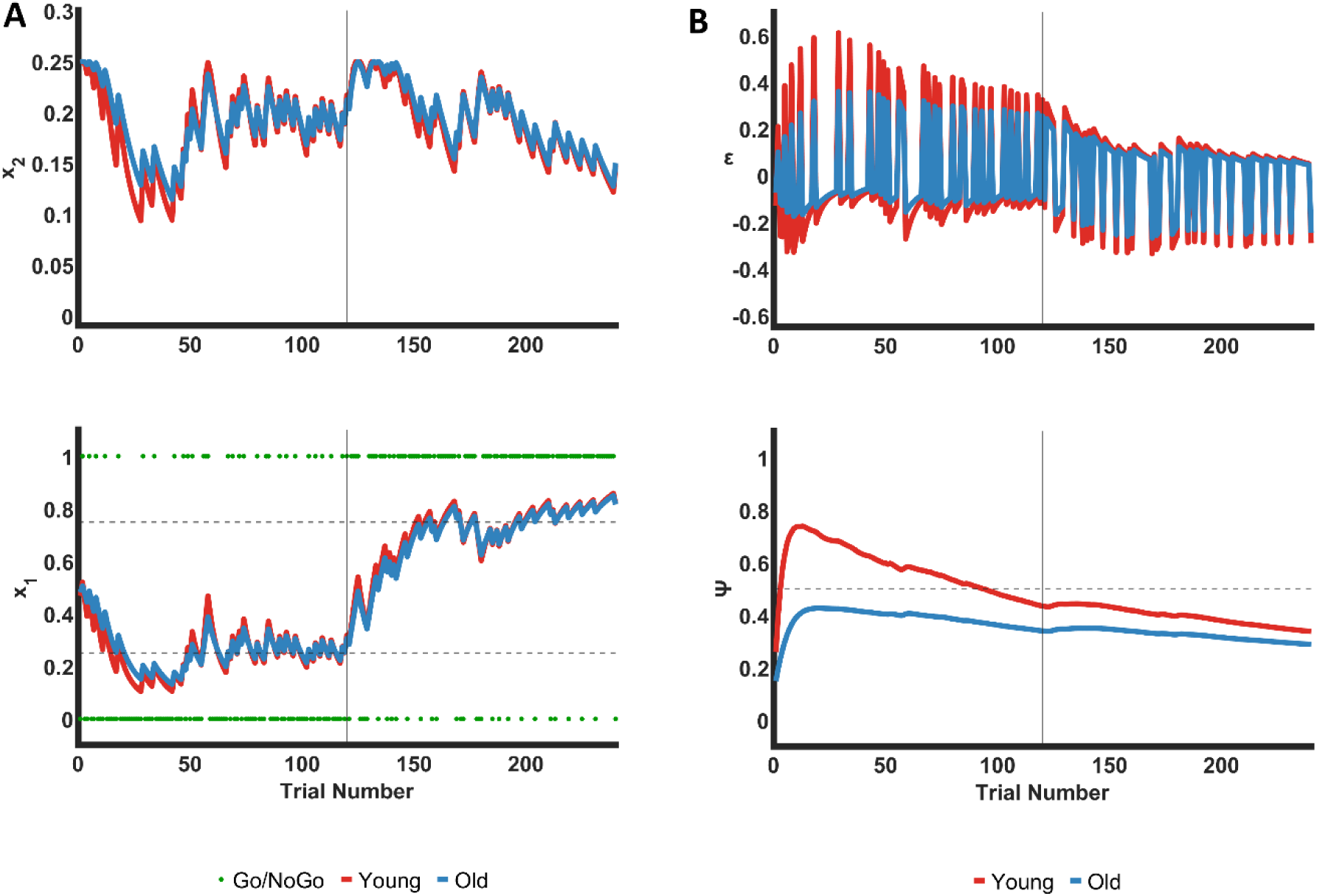
Simulated Hidden States in Perceptual Model of HGF. **(A)** Simulated posterior expectation (x_1_) and estimates of environmental uncertainty connected to β_4_ (x_2_) using the average parameter values of ω and β_4_ from young (red) and old (blue). The vertical solid line indicates the change point of condition (from mostly-no-go to mostly-go) and the horizontal dashed lines indicate expectancy of go-cue - 0.25 (25% go) and 0.75 (75% go). The green dots indicate binary inputs for no-go cue (0) and go cue (1) **(B)** Simulated precision-weighted prediction errors (ε) and precision weights (ψ) of prediction errors across trials in young (red) and old (blue). The horizontal dotted line indicates ψ = 0.5 and the vertical solid lines indicate the change point of condition (from mostly-no-go to mostly-go).

