## Supplementary Information for "Reduced belief updating impairs adaptive step initiation in older adults"

Jacob, A.B. *et al.*

**Table S1.** Descriptive statistics of the number of Go trials included in the analysis

| Go Trials | Group | Mean | Std. Deviation | Minimum | Maximum |
|---|---|---|---|---|---|
| Go Trials Left | young | 58.3 | 2.6 | 51.0 | 60.0 |
| Go Trials Left | old | 38.0 | 2.2 | 33.0 | 40.0 |
| Go Trials Right | young | 58.0 | 3.5 | 47.0 | 60.0 |
| Go Trials Right | old | 38.2 | 2.7 | 28.0 | 40.0 |
| Go Trials | young | 116.2 | 5.8 | 98.0 | 120.0 |
| Go Trials | old | 76.2 | 4.3 | 61.0 | 80.0 |

#### S1. Normalized Vertical Force

Vertical ground reaction force of stepping leg and stance leg were normalized to body weight (BW). BW can be estimated as the force of both legs averaged over time.

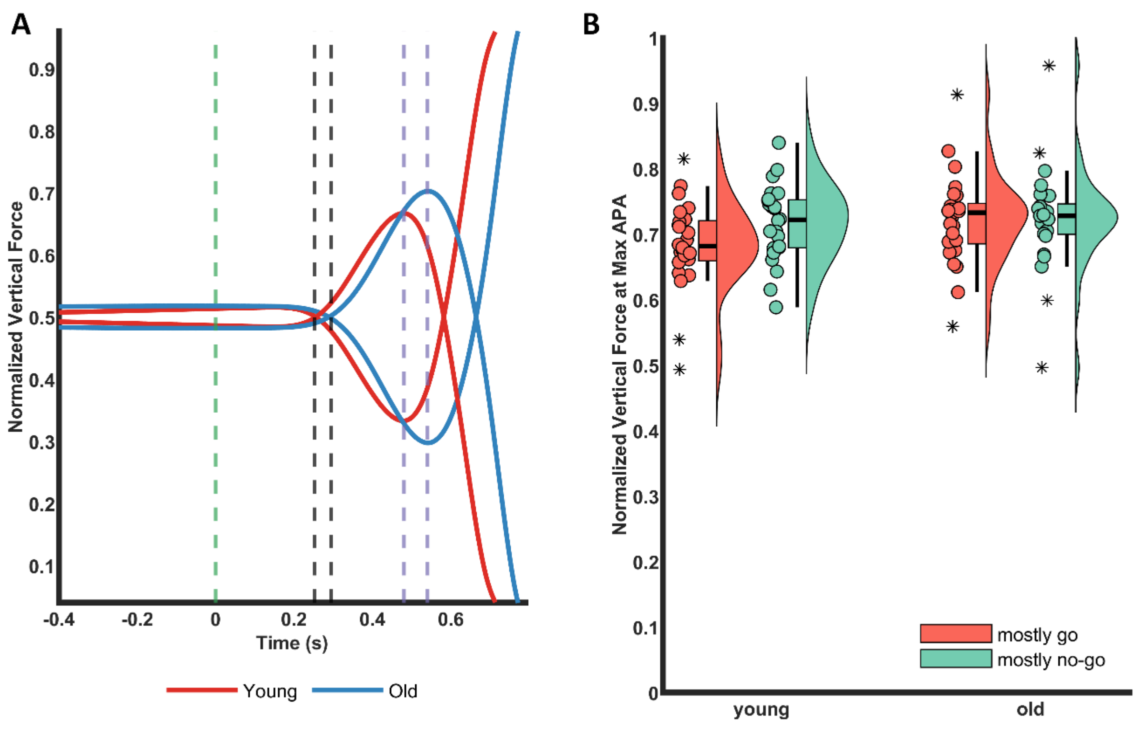


**Fig. S1.1 Force Averages (A)** Average vertical forces of young and old normalized to body weight. The green line indicates go cue, black lines indicate APA onsets, and purple lines indicate maximum APAs. On average, older people

show a higher maximum APA than younger people. **(B)** Raincloud plots of normalized vertical force of stepping leg at maximum APA as a function of condition (mostly go / mostly no-go) and groups (young/old).

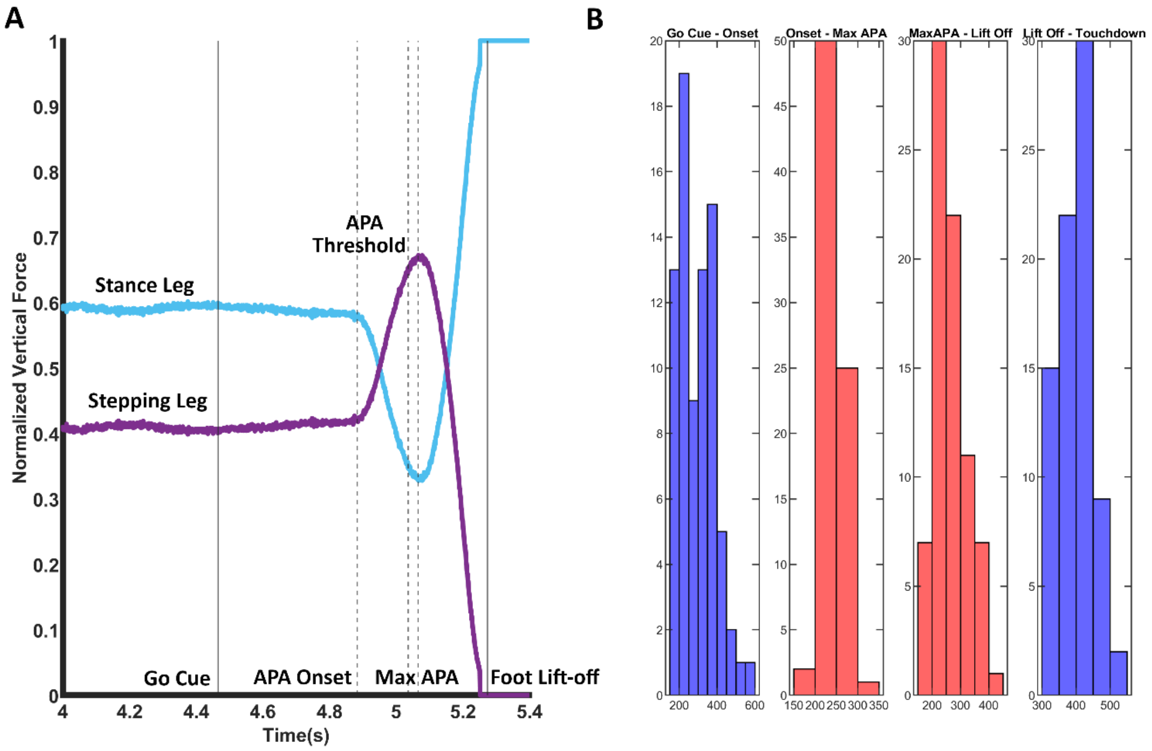


**Fig. S1.2 Maximum APA Detection (A)** Normalized vertical forces of stepping leg and stance leg in a go trial of an example participant. Here, the APA threshold was used to identify the maximum APA and APA onset. Maximum APA was identified as peak vertical force of stepping leg after the threshold and before foot lift-off. APA onset was identified as the first zero crossing point of the derivative before threshold **(B)** Histogram of a participant showing the distribution of latencies from go cue to APA onset, APA onset to maximum APA, maximum APA to foot lift-off, and foot lift-off to touchdown. The latencies of force trials present in the tail ends of the distribution were visualized to see if they were outliers or not.

**Table S1.1.** ANOVA Summary of Linear Mixed Model on Normalized Vertical Force of Stepping Leg at Maximum APA

| Effect | df | F | p |
|---|---|---|---|
| Condition | 1, 47.9 | 11.2 | 0.002 |
| Group | 1, 46.9 | 1.4 | 0.246 |
| Condition ✻ Group | 1, 47.9 | 10.9 | 0.002 |

**Table S1.2.** Contrasts of Linear Mixed Model on Normalized Vertical Force of Stepping Leg at Maximum APA

| | Estimate | SE | df | 95% CI | | z | p† |
|---|---|---|---|---|---|---|---|
| | | | | Lower | Upper | | |
| Mostly-go young vs old | -0.04 | 0.02 | ∞ | -0.08 | 0.00 | -1.98 | 0.190 |
| Mostly-no-go young vs old | -0.01 | 0.02 | ∞ | -0.05 | 0.03 | -0.30 | 1.000 |
| Young mostly-go vs mostly-no-go | 0.03 | 0.01 | ∞ | 0.02 | 0.05 | 4.26 | <.001 |
| Old mostly-go vs mostly-no-go | 0 | 0.01 | ∞ | -0.01 | 0.01 | 0.03 | 1.000 |

† p-values are adjusted using Bonferroni adjustment.

### S2. Force Latencies and Trial History

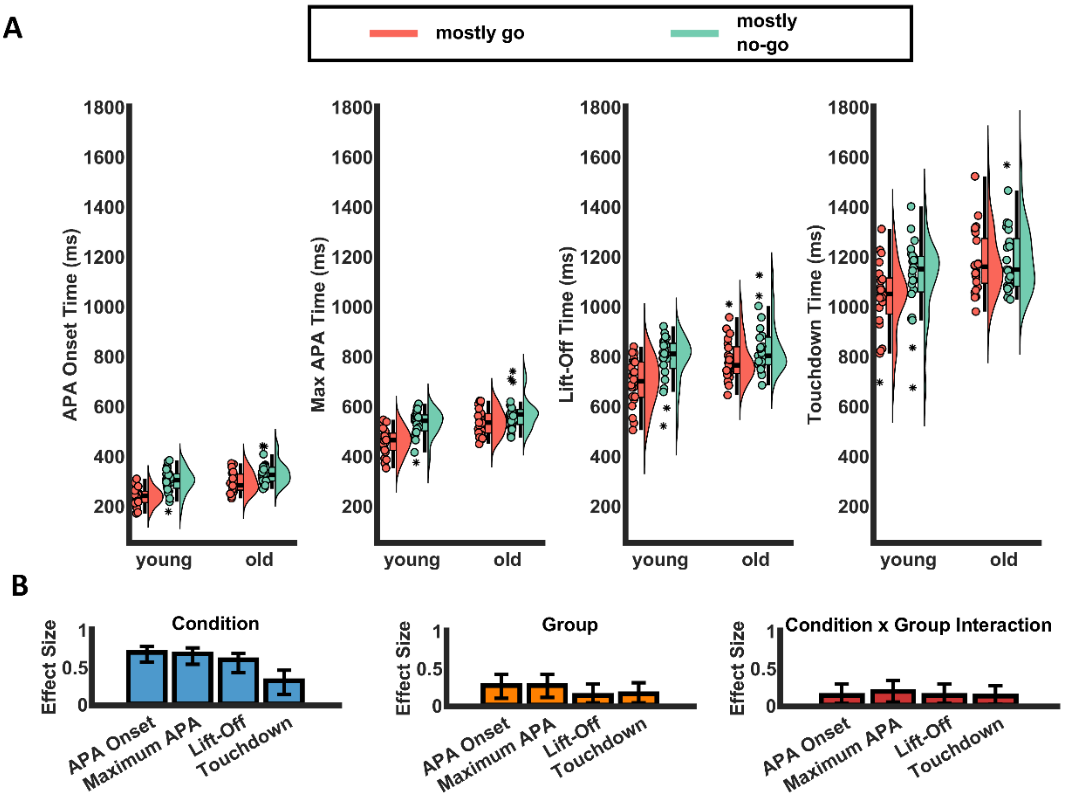


**Fig. S2.1 Reaction Time Descriptive Plots (A)** Raincloud plot of APA onset time, Maximum APA time, Lift-off time and Touchdown time of young and old participants. The scatter plot indicates individual time points, colours indicate conditions, and the whiskers indicate outliers **(B)** Bar plot of the estimated effect sizes ($\eta^2$ partial) of condition, group, and interaction for each of the four time points. Error bars indicate the 90% confidence interval.

**Table S2.1.** ANOVA Summary of Linear Mixed Model of APA Onset Time, Maximum APA Time, Lift-Off Time, Touchdown Time and Movement Time

| Dependent Variable | Effect | df | F | p | Partial $\eta^2$ | 90% CI Partial $\eta^2$ Lower | 90% CI Partial $\eta^2$ Upper |
|---|---|---|---|---|---|---|---|
| APA Onset Time | Condition | 1, 46.5 | 105.8 | < .001 | 0.69 | 0.56 | 0.77 |
| | Group | 1, 47.0 | 16.2 | < .001 | 0.26 | 0.09 | 0.41 |
| | Condition ✻ Group | 1, 46.5 | 7.0 | 0.011 | 0.13 | 0.02 | 0.28 |
| Maximum APA Time | Condition | 1, 46.6 | 94.2 | < .001 | 0.67 | 0.53 | 0.75 |
| | Group | 1, 47.0 | 16.2 | < .001 | 0.26 | 0.10 | 0.41 |
| | Condition ✻ Group | 1, 46.6 | 9.9 | 0.003 | 0.18 | 0.04 | 0.33 |
| Lift-Off Time | Condition | 1, 47.0 | 66.3 | < .001 | 0.59 | 0.42 | 0.68 |
| | Group | 1, 47.0 | 6.9 | 0.012 | 0.13 | 0.02 | 0.28 |
| | Condition ✻ Group | 1, 47.0 | 6.9 | 0.012 | 0.13 | 0.02 | 0.28 |
| Touchdown Time | Condition | 1, 47.0 | 20.8 | < .001 | 0.31 | 0.13 | 0.45 |
| | Group | 1, 47.0 | 8.4 | 0.006 | 0.15 | 0.03 | 0.30 |
| | Condition ✻ Group | 1, 47.0 | 6.2 | 0.016 | 0.12 | 0.01 | 0.26 |
| Movement Time | Condition | 1, 44.3 | 10.8 | 0.002 | 0.19 | 0.05 | 0.34 |
| | Group | 1, 44.1 | 4.8 | 0.033 | 0.09 | 0.00 | 0.23 |
| | Condition ✻ Group | 1, 44.3 | 1.4 | 0.245 | 0.03 | 0.00 | 0.14 |

**Table S2.2.** Contrasts of Linear Mixed Model on APA onset time

| | | | | 95% CI | | | |
|---|---|---|---|---|---|---|---|
| | Estimate | SE | df | Lower | Upper | z | p† |
| mostly go young vs mostly go old | 58.0 | 11.1 | ∞ | 36.2 | 79.7 | -5.23 | < .001 |
| mostly no-go young vs mostly no-go old | 34.7 | 14.4 | ∞ | 6.5 | 62.9 | -2.41 | 0.064 |
| mostly go young vs mostly no-go young | -60.93 | 7.7 | ∞ | -52.8 | -22.5 | -4.86 | < .001 |
| mostly go old vs mostly no-go old | -37.65 | 12.6 | ∞ | -75.18 | -46.74 | -8.40 | < .001 |

† p-values are adjusted using Bonferroni adjustment.

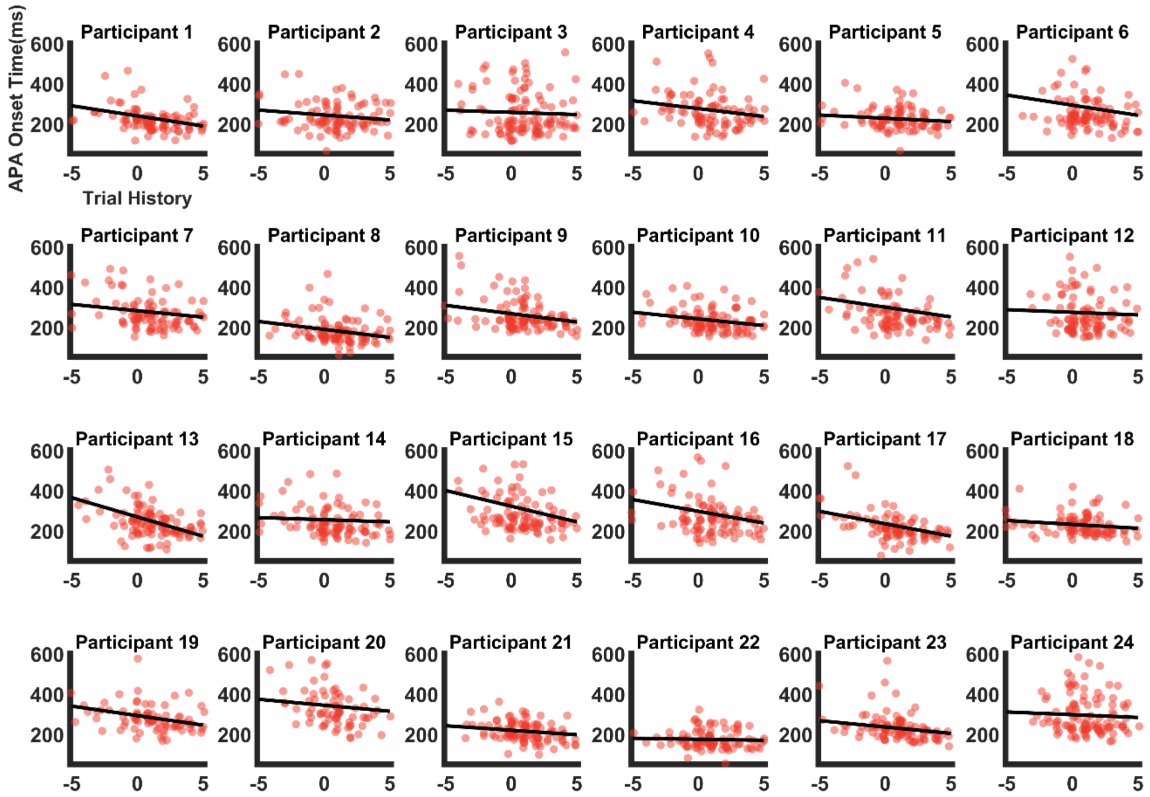


**Fig. S2.2** Scatter plots of the APA onset times of individual trials a function of trial history of the young participants

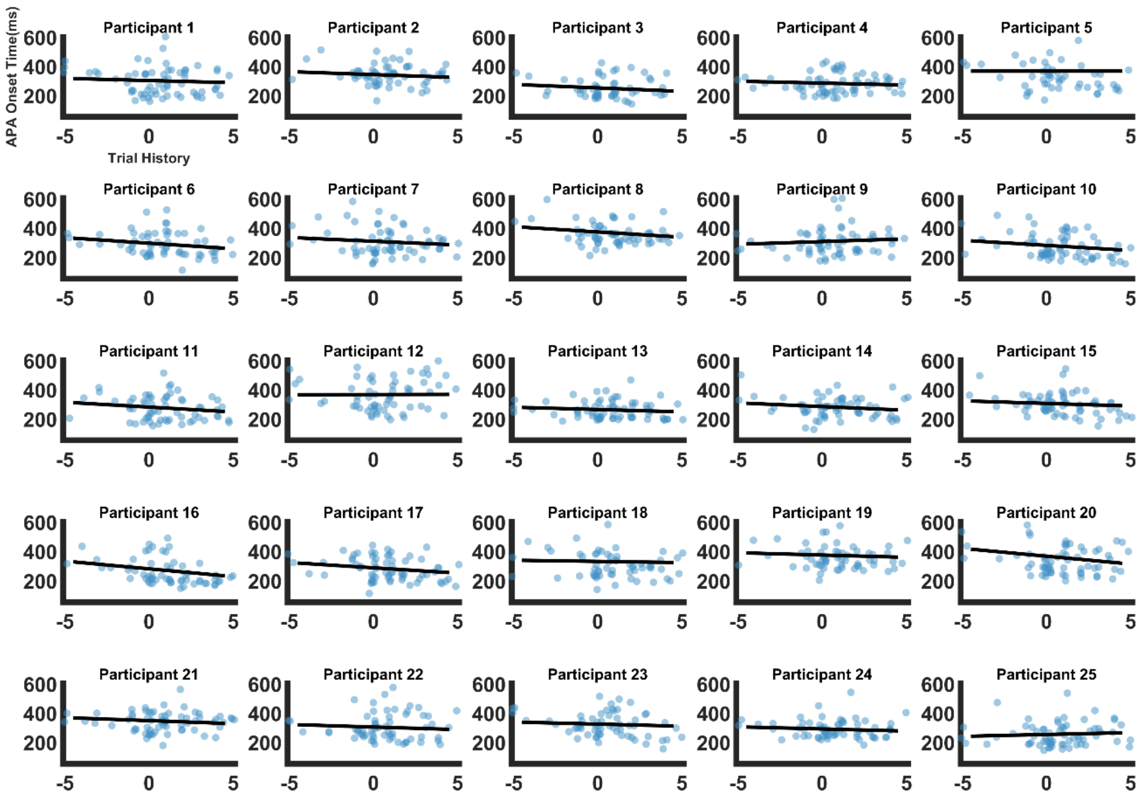


**Fig. S2.3** Scatter plots of the APA onset times of individual trials a function of trial history of the old participants

**Table S2.3.** ANOVA Summary of Linear Mixed Model on Trial History on APA Onset Time, Maximum APA Time, Lift-Off Time, and Touchdown Time

| Dependent Variable | Effect | df | F | p | Partial $\eta^2$ | 90% CI Partial $\eta^2$ Lower | 90% CI Partial $\eta^2$ Upper |
|---|---|---|---|---|---|---|---|
| APA Onset Time | Trial History | 1, 43.4 | 101.7 | < .001 | 0.70 | 0.56 | 0.77 |
| | Group | 1, 47.0 | 19.1 | < .001 | 0.29 | 0.12 | 0.44 |
| | Trial History ✻ Group | 1, 43.4 | 7.7 | 0.008 | 0.15 | 0.02 | 0.31 |
| Maximum APA Time | Trial History | 1, 45.0 | 122.3 | < .001 | 0.73 | 0.61 | 0.80 |
| | Group | 1, 47.0 | 19.1 | < .001 | 0.29 | 0.12 | 0.44 |
| | Trial History ✻ Group | 1, 45.0 | 12.0 | 0.001 | 0.21 | 0.06 | 0.37 |
| Lift-Off Time | Trial History | 1, 46.0 | 101.9 | < .001 | 0.69 | 0.55 | 0.76 |
| | Group | 1, 47.0 | 8.1 | 0.006 | 0.15 | 0.03 | 0.30 |
| | Trial History ✻ Group | 1, 46.0 | 8.9 | 0.005 | 0.16 | 0.03 | 0.31 |
| Touchdown Time | Trial History | 1, 45.1 | 54.7 | < .001 | 0.55 | 0.37 | 0.66 |
| | Group | 1, 47.1 | 9.8 | 0.003 | 0.17 | 0.04 | 0.32 |
| | Trial History ✻ Group | 1, 45.1 | 6.6 | 0.014 | 0.13 | 0.02 | 0.28 |

### S3. Hierarchical Gaussian Filter

The response model used in our study is a log-reaction time model (Log (RT)). Log (RT) is described as a function of six gaussian parameters: $\beta_0$, $\beta_1$, $\beta_2$, $\beta_3$, $\beta_4$, and ζ. These coefficients of the response model describe how trial-wise beliefs are translated into log-reaction times. The parameters $\beta_{0\text{-}4}$ capture the weights of surprise of the stimulus outcome (Go/No-Go), certainty of the outcome, probability of informational uncertainty, environmental uncertainty, and Gaussian decision noise on the predicted log (RT). The default equation of the Log RT model is given as follows:

$$\log(\mathrm{RT})^{t} = \beta_0 + \beta_1 * (\delta_1)^{t} + \beta_2 * (\varepsilon_3)^{t} + \beta_3 * (\mu_3)^{t} + \beta_4 * (p.v)^{t} + \zeta^{t}$$

(Eq. S3.1)

$$p.v = \sigma(\mu_2).[1 - \sigma(\mu_2)]e^{\mu_3}$$

(Eq. S3.2)

In Eq. S3.1, $\delta_1$ is surprise, $\varepsilon_3$ is Bernoulli variance, $\mu_3$ is inferential variance, and $p.v$ is phasic volatility (Eq. S3.2). Because the task structure and RT distribution deviated from canonical paradigms previously fitted with HGF, we first calibrated the model by performing a prior predictive check of the HGF linear response model for log RTs. Specifically, we simulated RTs across a range of response model parameters while keeping the perceptual model fixed to Bayes-optimal values. We compared the simulated RT distributions to the empirical RT density averaged across participants, adjusting the response model's prior means until the simulated distributions qualitatively reproduced the empirical RT distribution. These tuned priors were then used as group-level priors in subsequent model fitting and between-group comparisons. The resulting prior parameter values were as follows: $\beta_0$ ($\mu$ = log (200); $\sigma$ = 4), $\beta_1$ ($\mu$ = 0; $\sigma$ = 4), $\beta_2$ ($\mu$ = -2; $\sigma$ = 4), $\beta_3$ ($\mu$ = 2; $\sigma$ = 4), $\beta_4$ ($\mu$ = 2; $\sigma$ = 4), and ζ ($\mu$ = log (log (20)); $\sigma$ = 4).

The tonic learning rate ω is a function of outcome uncertainty, informational uncertainty, and environmental uncertainty. The equation of learning rate ω is given as follows:

$$\omega = \frac{\hat{\pi}_u}{\dfrac{1}{\sigma_1^{(k-1)} + e.v} + \hat{\pi}_u}$$

(Eq. S3.3)

$$e.v = e^{(\kappa_1 \mu_2^{(k-1)} + \omega_1)}$$

(Eq. S3.4)

In Eq. S3.3, $\hat{\pi}_u$ is the outcome uncertainty, $\sigma_1^{(k-1)}$ is the informational uncertainty, and $e.v$ (Eq. S3.4) is the environmental uncertainty.

**Table S3.1.** Independent samples t-test of perceptual and response model parameters

| | t | df | p | p† corrected | Cohen's d | SE Cohen's d | 95% CI Cohen's d Lower | 95% CI Cohen's d Upper |
|---|---|---|---|---|---|---|---|---|
| ω | 3.891 | 47 | <.001 | 0.002 | 1.11 | 0.33 | 0.50 | 1.71 |
| $\beta_0$ | -3.194 | 47 | 0.003 | 0.009 | -0.91 | 0.32 | -1.50 | -0.32 |
| $\beta_1$ | 1.631 | 47 | 0.109 | 0.159 | 0.47 | 0.30 | -0.10 | 1.03 |
| $\beta_2$ | 2.090 | 47 | 0.042 | 0.080 | 0.60 | 0.30 | 0.02 | 1.17 |
| $\beta_3$ | -0.329 | 47 | 0.744 | 0.868 | -0.09 | 0.29 | -0.65 | 0.47 |
| $\beta_4$ | -2.675 | 47 | 0.010 | 0.027 | -0.76 | 0.31 | -1.34 | -0.18 |
| ζ | 0.133 | 47 | 0.895 | 0.895 | 0.04 | 0.29 | -0.52 | 0.60 |

**Table S3.2.** Descriptive Statistics of perceptual and response model parameters

| | Group | N | Mean | SD | SE | Coefficient of variation |
|---|---|---|---|---|---|---|
| ω | young | 24 | -2.73 | 1.04 | 0.21 | -0.38 |
| | old | 25 | -3.91 | 1.08 | 0.22 | -0.28 |
| $\beta_0$ | young | 24 | 5.40 | 0.28 | 0.06 | 0.05 |
| | old | 25 | 5.63 | 0.28 | 0.06 | 0.05 |
| $\beta_1$ | young | 24 | 0.11 | 0.09 | 0.02 | 0.83 |
| | old | 25 | 0.08 | 0.06 | 0.01 | 0.81 |
| $\beta_2$ | young | 24 | -1.22 | 1.39 | 0.29 | -1.14 |
| | old | 25 | -1.90 | 0.80 | 0.16 | -0.42 |

**Table S3.2.** Descriptive Statistics of perceptual and response model parameters

| | Group | N | Mean | SD | SE | Coefficient of variation |
|---|---|---|---|---|---|---|
| $\beta_3$ | young | 24 | 1.80 | 0.88 | 0.18 | 0.49 |
| | old | 25 | 1.87 | 0.79 | 0.16 | 0.42 |
| $\beta_4$ | young | 24 | 0.12 | 0.70 | 0.14 | 5.72 |
| | old | 25 | 0.57 | 0.46 | 0.09 | 0.81 |
| ζ | young | 24 | 0.07 | 0.03 | 0.01 | 0.39 |
| | old | 25 | 0.07 | 0.03 | 0.01 | 0.39 |

† p-values are adjusted using Benjamini-Hochberg correction

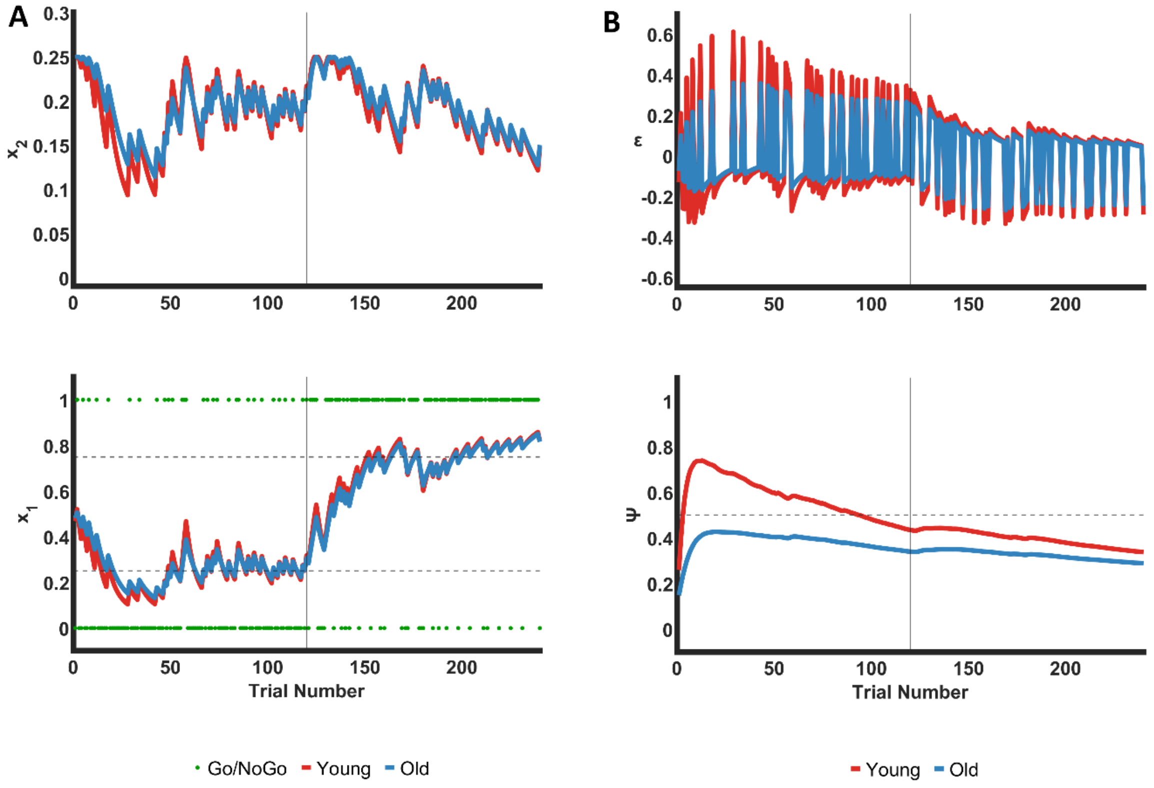


**Fig. S3.1 Simulated Hidden States in Perceptual Model of HGF (A)** Simulated posterior expectation ($x_1$) and estimates of environmental uncertainty connected to $\beta_4$ ($x_2$) using the average parameter values of ω and $\beta_4$ from young (red) and old (blue). The vertical solid line indicates the change point of condition (from mostly-no-go to mostly-go) and the horizontal dashed lines indicate expectancy of go-cue - 0.25 (25% go) and 0.75 (75% go). The green dots indicate binary inputs for no-go cue (0) and go cue (1) **(B)** Simulated precision-weighted prediction errors (ε) and precision weights (ψ) of prediction errors across trials in young (red) and old (blue). The horizontal dotted line indicates ψ = 0.5 and the vertical solid lines indicate the change point of condition (from mostly-no-go to mostly-go).
